# Protrudin acts at ER-endosome contacts to promote KIF5-mediated endosomal fission and endosome-to-Golgi transport

**DOI:** 10.1101/2024.07.15.602703

**Authors:** Julia Kleniuk, Aishwarya G. Nadadhur, Emily Wolfenden, Catherine Rodger, Eliska Zlamalova, Evan Reid

## Abstract

Fission of transport tubules from early endosomes is required for endosomal sorting, but mechanisms of endosomal tubule fission (ETF) are incompletely understood. We show protrudin acts at ER-endosome contacts to promote ETF and endosome-to-Golgi traffic. Protrudin-mediated ETF required its ability to interact with ER-localised VAP proteins, endosomal phosphoinositides and KIF5. These properties also regulated the distance between protrudin and endosomal tubules. The defective ETF phenotype of increased endosomal tubulation in cells lacking protrudin was phenocopied by depletion of KIF5, but not FYCO1, a motor protein adaptor implicated in protrudin-dependent late endosome motility. It also required intact microtubules and dynein, consistent with a model where protrudin facilitates a tug-of-war between KIF5 and dynein to fission tubules. In addition to its direct role, protrudin links many other machineries involved in ETF, thus our findings elucidate how ETF is co-ordinated. These machineries are enriched for proteins implicated in hereditary motor neuron disorders, and protrudin or KIF5 depletion caused defective ETF in human neurons.

**Summary:** Protrudin binds ER-localised VAPs and endosomal phosphoinositides to form ER-endosome contacts that promote endosomal tubule fission and endosome-to-Golgi traffic. Protrudin recruits KIF5 to provide a FYCO1-independent force to fission endosomal tubules in neurons and non-polarised cells.

## Introduction

The early sorting endosome (ESE) is a key intracellular trafficking hub. Sorting decisions in this compartment control processes including cell signalling, cell-cell interactions, cell metabolism, cell motility and nutrient uptake, as well as protein and organelle degradation.[1, 2] In addition to being physiologically necessary, these processes impact on human diseases, from the very common, such as cancer, atherosclerosis, metabolic disorders and Alzheimer’s disease, to rare but devastating neurogenetic conditions.[3–6] It is therefore crucial that the molecular mechanisms underlying early endosomal sorting are fully understood.

In the ESE, cargoes that have been endocytosed from the plasma membrane are either sorted away from the endosomal system to destinations such as the plasma membrane or Golgi apparatus, or retained for traffic to the late endosomal-lysosomal degradative compartment.[1, 2] Traffic away from endosomes involves sorting of cargoes into endosomal tubules (ETs), which form from the endosomal body, elongate and then fission.[1, 2] Subtypes of ETs are specialised for traffic of specific cargoes, and these are differentially marked by members of the sorting nexin (SNX) family of proteins, for example, SNX1 in the case of tubules that traffic from endosomes to the Golgi apparatus.[7, 8]

Fission from the parent endosome is necessary for forward transport of ETs and the complex molecular mechanisms that drive this process are beginning to be understood. A microtubule-dependent pulling force is important, with early studies showing that the long axis of ETs aligns along microtubules and that ET fission (ETF) requires oppositely-directed forces provided by kinesin (especially KIF5 and KIF13A) and dynein microtubule motor proteins.[9–13] Machineries that directly promote membrane deformation are also involved. A helical complex comprised of the atypical endosomal sorting complex required for transport-III (ESCRT-III) proteins IST1 (increased sodium tolerance 1) and CHMP1B (charged multivesicular body protein 1B) promotes ETF by constricting tubules from the outside.[12, 14] In vitro reconstitution studies suggest that while not sufficient in itself to fission tubules, when combined with a pulling force on the tubule fission can be achieved.[15] Enzymatic proteins, such as dynamin GTPases or the ATPase EHD1 (EPS15 homology domain-containing protein 1), can also assemble on the tubule to drive membrane constriction.[16–18] In addition, efficient fission requires assembly of a branched actin network on the endosomal body. This is achieved by the WASH (Wiskott Aldrich Syndrome protein and scar homologue) complex, which activates the branched actin nucleator ARP2/3 (actin related protein complex 2/3) to achieve this.[19, 20]

Recently it has been recognised that many of the mechanisms involved in ETF are promoted and co-ordinated by membrane contact sites (MCS) between endosomes and the endoplasmic reticulum (ER).[21] The site of membrane breakage during ETF is at contacts between ETs and ER tubules, in a process termed “ER-associated” ETF (ER-ETF), and correct ER morphogenesis is required for this process to occur. Our studies on the microtubule modelling ATPase spastin and IST1 provided the first description of the molecular machinery at ER-endosome MCS involved in ER-ETF. We showed that interaction between endosomal IST1 (see above) and the ER-localised microtubule severing enzyme spastin is required for efficient fission of SNX1-positive ETs.[12, 22] Spastin presumably acts in this process to modify microtubules to promote motor-dependent force activity along the breaking ET.

Since then, several other molecular mechanisms that regulate ER-ETF have been elucidated. Vesicle associated membrane protein-associated proteins A and B (VAPA/B), which participate in many functionally important interactions at MCS between the ER and other organelles, play a critical role. At ER-endosome contacts the VAPs interact with and promote the activity of several lipid transfer proteins. These include oxysterol binding protein (OSBP), a lipid transporter that transfers phosphatidylinositol-4-phosphate (PI4P) from the endosome to the ER, and oxysterol binding protein-related proteins 9 and 10 (ORP9 and ORP10), which regulate endosomal PI4P concentrations by counter-transport of PI4P and phosphatidylserine (PS) between endosomes and the ER. In cell lacking VAPs or OSBP there is increased endosomal PI4P, accompanied by ectopic WASH complex-mediated actin formation on endosomes and blocked traffic of endosome-to-Golgi cargoes, while ORP9/10-mediated PS insertion into the endosomal membrane promotes ETF by increasing recruitment of the membrane scission protein EHD1.[23, 24] The branched actin network on endosomes is also regulated by an interaction between the ER-localised TMCC1-3 (transmembrane and coiled-coil domain family 1-3) proteins and endosomal coronin 1 proteins. The coronins act to disassemble Arp2/3-containing branched actin filaments and so restrict actin from the ET. Cellular depletion of coronins allows actin to spread along the ET, preventing formation of the ER-ET contacts that are critical for ETF and cargo sorting.[25, 26] It is not entirely clear how all of the disparate mechanisms involved in ETF are co-ordinated, although coronin 2a binds to EHD1, linking actin-related mechanisms in ETF to another membrane fission machinery.[27]

The importance of elucidating the mechanisms of ETF is exemplified by its role in the pathogenesis of hereditary motor neuron disorders, especially hereditary spastic paraplegias (HSPs). HSPs are genetic neurological disorders in which the longest axons of the corticospinal tracts are affected by a dying back axonopathy, resulting in the development of progressive spastic paralysis of the legs.[28] To date, four proteins encoded by genes mutated in HSP have been implicated in ETF; spastin, strumpellin, atlastin-1 and dynamin 2. Mutations in the gene encoding spastin (see above) are the commonest cause of autosomal dominant HSP in northern Europe and North America, while mutations in WASHC5 cause a relatively rare form of autosomal dominant HSP.[29, 30] WASHC5 encodes strumpellin, a member of the WASH complex and cells lacking strumpellin develop increased endosomal tubulation, consistent with a defect in ETF.[31] Recently we have described an endosomal tubulation phenotype in neurons lacking atlastin-1, mutations in which are one of the most common causes of childhood onset autosomal dominant HSP.[32] Finally, mutations in the gene encoding the membrane scission protein dynamin 2 cause a range of neuronal disorders, ranging from hereditary motor and sensory neuropathy to HSP.[33, 34]

At least four proteins involved in ER-ETF, the VAPs, spastin, atlastin-1 and OSBP, interact with protrudin, an ER-localised integral membrane protein involved in ER-endosome MCS.[35, 36] Protrudin was named after its ability to induce long cellular protrusions in normally non-polarised cell types and to promote neurite extension in neurons.[37] More recently protrudin has been demonstrated to be a potent enhancer of central nervous system axonal regeneration after injury.[38] It is anchored in the ER by N-terminal hydrophobic domains (HDs), and it interacts with ER-localised VAPs via a FFAT (two phenylalanines in an acidic tract) motif (Figure 1A).[36, 37, 39] Protrudin forms ER-late endosome MCS via coincidence detection involving binding to endosomal RAB7 and, via a FYVE (Fab 1, YOTB, Vac 1, and EEA1) domain, to phosphatidylinositol-3-phosphate (PI3P).[40] Protrudin also interacts with the kinesin motor KIF5 and at the ER-late endosome MCS it transfers KIF5 to the endosomal motor protein adaptor FYCO1 (FYVE and coiled-coil domain autophagy adaptor 1).[40, 41] This promotes microtubule plus-end directed polarised traffic of late endosomes towards the tips of protrusions, where they fuse with the plasma membrane to deliver membrane for protrusion extension. Protrudin also has a RAB11 binding domain, which preferentially binds to RAB11-GDP, locking RAB11 in this inactive state. As the RAB11 compartment is involved in recycling cargoes to the plasma membrane, this has been proposed to switch off generalised (i.e. non-polarised) recycling.[37]

**Figure 1.**
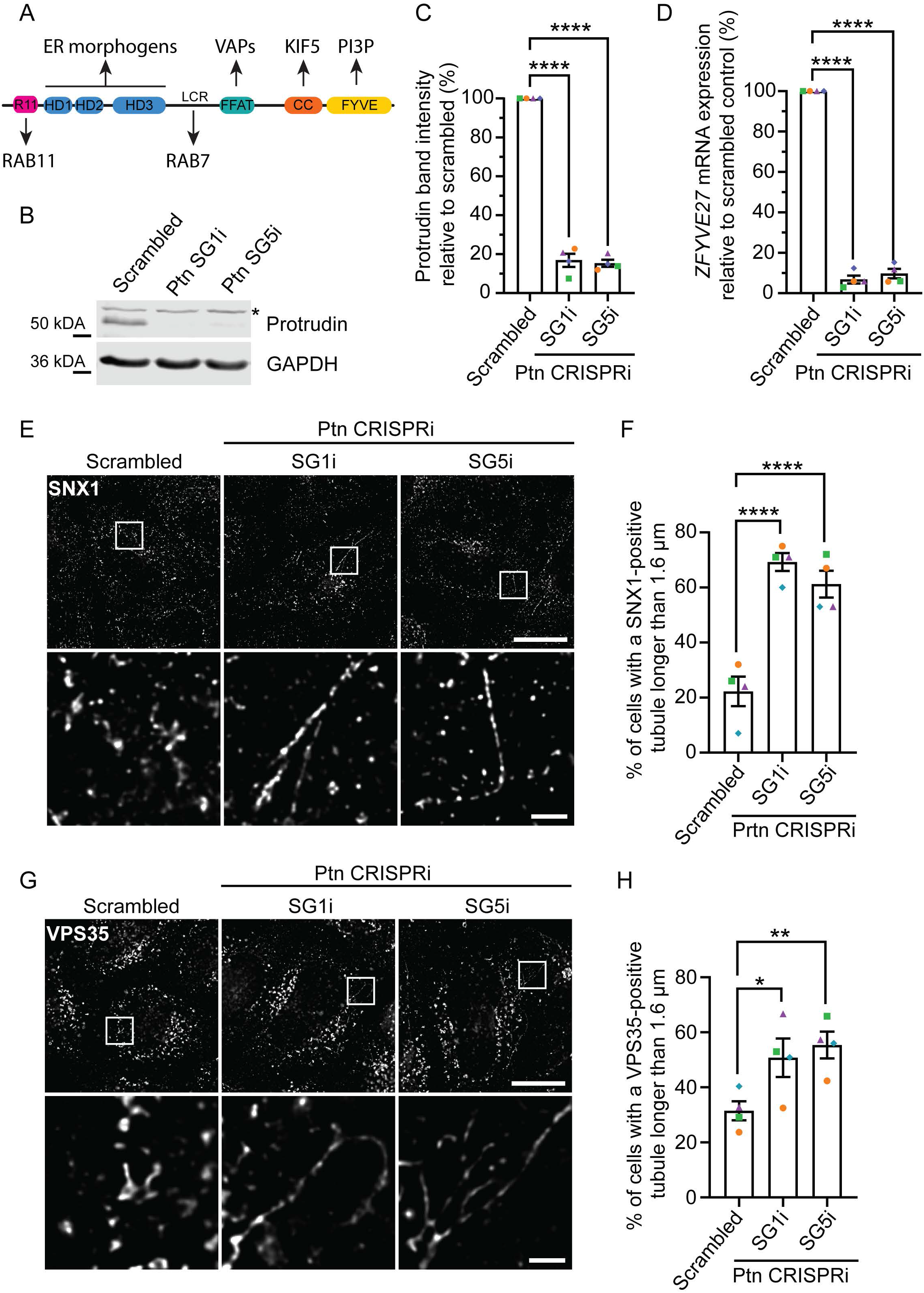
HeLa cells depleted of protrudin have increased endosomal tubulation. **A)** Cartoon of the domain structure of protrudin. R11 - Rab11 binding domain; HD - hydrophobic domain; LCR – low complexity region, FFAT - two phenylalanines in an acidic tract motif; CC – coiled-coil domain; FYVE - FYVE domain. **B**) HeLa cells stably expressing CRISRPi machinery were transduced with a scrambled control single guide (SG) RNA or SG RNAs targeting the protrudin (Ptn) transcriptional start site (SG1i and SG5i). Cells were lysed and immunoblotted against protrudin. GAPDH serves to verify equal lane loading. * - non- specific band. Protrudin band intensity was normalised to GAPDH band intensity and the resulting values expressed as a percentage of the scrambled control value. Quantification of 4 such experiments is plotted in (**C**). **D**) *ZFYVE27* (the gene encoding protrudin) transcript abundance was measured by RT-qPCR in the cell lines indicated, and the results for 4 biological repeats are shown. *ACTB* was used as a housekeeping control to normalise the results and mRNA expression in the protrudin CRISPRi lines was expressed as percentage of the scrambled control. **E**) The HeLa lines indicated were fixed and labelled for endogenous SNX1. The lower panel images are magnified views of the boxed areas in the corresponding upper panel image. **F**) Quantification of **E**). The percentage of cells with a SNX1-positive tubule longer than 1.6 μm was plotted. Data from 4 biological repeats are shown, 100 cells per condition per repeat analysed. Bars show mean +/- S.E.M. **G**) The cell lines indicated were fixed and labelled for endogenous VPS35 and the percentage of cells with a long tubule was quantified in 4 biological repeats as described for (**F**), except 103-122 cells per condition per repeat were analysed. Statistical testing was done by repeated measures one-way ANOVA with Dunnett correction for multiple comparisons. Micrographs scale bars = 20 μm, inset scale bars = 2 μm.

Considering protrudin’s participation in ER-endosome contacts and its interaction with several other proteins implicated in ER-ETF, in this paper we investigate whether protrudin has a role in this process. We found that protrudin depletion resulted in an increased number of long ETs, inefficient ETF and inefficient endosome-to-Golgi traffic of the cation independent mannose 6-phosphate receptor (ciM6PR), a receptor that traffics through endosomal tubules. Rescue experiments demonstrated that protrudin required the ability to bind to VAPs, KIF5 and phosphatidylinositol, but not the ability to bind to RAB11 or insert into the ER membrane, to support ETF and ciM6PR traffic. Super-resolution microscopy experiments showed that protrudin colocalised with ET markers, but the physical distance between protrudin and endosomes was altered in the mutants that failed to rescue ETF and trafficking. The elongated ETs that developed in the absence of protrudin required the presence of intact and dynamic microtubules and functional dynein. A similar endosomal tubulation phenotype was also observed after depletion of KIF5 but not of FYCO1. Finally, we showed that these observations were relevant to human cortical neurons. Thus, we identify a new role for protrudin in regulation ER-associated ETF. Our results are consistent with a model where protrudin regulates this process by promoting a KIF5-dependent, FYCO1-independent force directed towards the plus ends of microtubules, which provides energy for fission when counterbalanced by a dynein-dependent force directed towards the microtubule minus-end. Furthermore, via its interactions with VAPs, KIF5, atlastin-1 and spastin, protrudin links ETF machineries involved in actin and phospholipid regulation, microtubule force generation, endosomal membrane deformation and ER shaping, and thus is a key factor in co-ordinating the processes involved in ETF.

## Results

### Protrudin regulates endosomal tubulation in HeLa cells

We first investigated whether protrudin depletion affects endosomal tubulation. A CRISPRi-mediated gene silencing approach was used to reduce protrudin expression in HeLa cells.

HeLa cells stably expressing CRISPRi machinery (KRAB transcriptional repressor fused to a dead Cas9 enzyme) were generated, allowing stable silencing of genes of interest upon lentiviral transduction of a single guide RNA (sgRNA) targeting the relevant transcriptional start site (Supplementary Figure 1A).[42] We tested multiple sgRNAs targeting the transcription start site of the human protrudin gene (*ZFYVE27*) and two guides with the best performance as measured by reduction in *ZFYVE27* mRNA expression were validated by immunoblotting and additional RT-qPCR, and used for further experiments (protrudin-SG1i and -SG5i; Supplementary Figure 1B, Figure 1B-D). *ZFYVE27* expression in these lines was monitored periodically and showed at least 80% depletion throughout the study.

Cellular depletion of spastin results in the formation of cellular protrusions that require protrudin for their formation, and so to validate our cell lines using a known functional effect of protrudin depletion, we tested the effects of spastin depletion on protrusion formation.[43] Consistent with previous results, we found that siRNA-mediated depletion of spastin induced cellular protrusions in control cells but not in protrudin-SG1i and -SG5i cells (Supplementary Figure 1C-E).

Next we examined the effects of protrudin depletion on ETs, using SNX1 and the retromer complex member vacuolar protein sorting 35 (VPS35) as markers. SNX1 associates with the retromer complex and together they localise to ETs responsible for retrograde trafficking of cargoes from endosomes to the Golgi apparatus.[7, 8] For both markers, we found that protrudin depletion caused an increased proportion of cells with long tubules, defined as being >1.6 μm in length (Figure 1E-H). Consistent with their known function, these two markers co-localised to a similar degree in control cells and in protrudin-depleted cells, and in protrudin-depleted cells co-localised domains were observed along the elongated ETs (Supplementary Figure 2). We concluded that protrudin is required for proper regulation of the length of ETs.

### Protrudin is required for endosomal tubule fission

Increased endosomal tubulation can be caused by defective ETF or by induction of tubule formation. To distinguish between these two possibilities, we carried out experiments to quantify the dynamics of ETF in living cells lacking protrudin. To do these experiments efficiently we used MRC5 human fibroblast cells, as we have previously found the flat morphology of these cells facilitates live cell imaging of ETs. To generate a suitable system for this work we first FACS sorted an existing GFP-SNX1 expressing MRC5 cell line to select low-to-medium expressing cells, to minimise over-expression artefacts (Figure 2A).[22] We then stably expressed CRISPRi machinery in the cells using lentiviral transduction, then further transduced the cells with scrambled and protrudin-SG1i and -SG5i guides to inhibit protrudin expression. We confirmed that each cell line was expressing GFP-SNX1 at the expected size and with equal abundance, and that protrudin was depleted in the lines where it was targeted (Figure 2B-C). We then analysed ET dynamics in these cells, and found that mean tubule duration, defined as the time taken from first appearance of a tubule until its fission, collapse back into the parent endosome (indicative of failed fission) or the end of the movie, was significantly increased in both cell lines depleted of protrudin (Figure 2D & E, Supplementary Movies 1-3). This strongly suggests that protrudin is required for efficient fission of ETs from the parent endosome.

**Figure 2.**
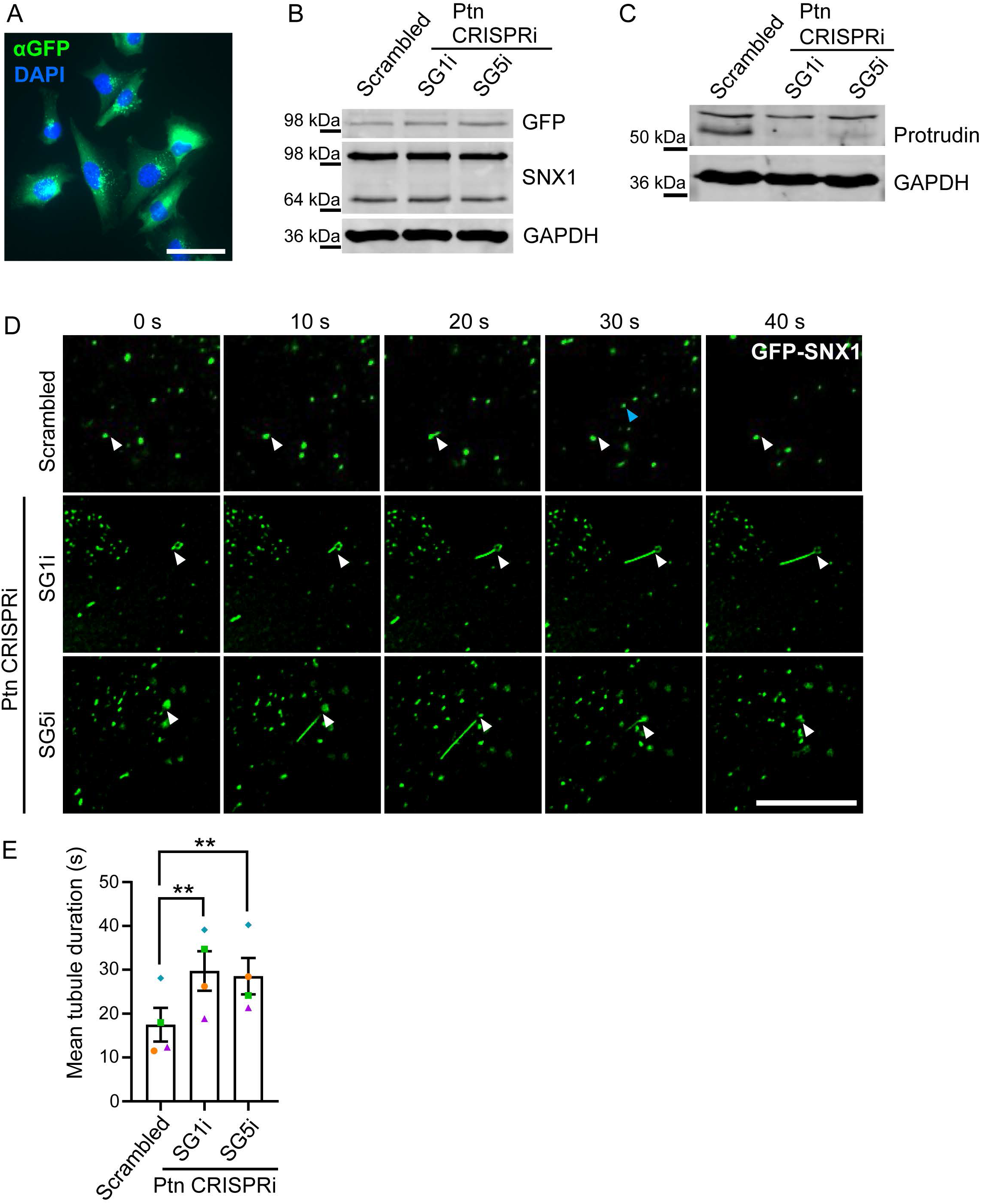
Protrudin depletion inhibits endosomal tubule fission. **A)** MRC5 fibroblasts stably expressing GFP-SNX1 were FACS sorted to select low-to-medium expressing cells then transduced to generate stable expression of CRISPRi machinery. Cells were fixed and stained with an anti-GFP antibody and DAPI. Scale bar = 50 μm. **B**) CRISPRi-GFP-SNX1 cells were transduced with SG RNAs shown, lysed and immunoblotted with the antibodies indicated. In the SNX1 immunoblot, the upper band is exogenously expressed GFP-SNX1 and lower band is endogenous SNX1. **C**) Protrudin depletion in cell lines expressing protrudin SG RNAs was validated by immunoblotting against endogenous protrudin. GAPDH labelling serves to validate equal lane loading. **D**) Representative stills from Supplementary Movies 1-3 showing GFP-SNX1-positive ETs in the CRISPRi-GFP-SNX1 lines indicated. White arrowheads point to the position from which a tubule arises from an endosome, blue arrowhead is a fissioned tubule. Movies were acquired using an AxioObserver widefield microscope for 85- 90 seconds at a frame rate of 500 ms. Scale bar = 10 μm. **E**) Quantification of the duration of GFP-SNX1 tubules shown in D). Mean +/- S.E.M. for 4 independent biological repeats is shown; 5 tubules in 5 cells per condition per repeat were analysed. Statistical testing was with repeated measures one-way ANOVA with Dunnett correction for multiple comparisons.

### Protrudin mediates efficient endosome-to-Golgi traffic

We next examined whether the defective ETF in cells lacking protrudin had consequences for cargo trafficking. Since we had observed defective fission of SNX1 and VPS35-labelled tubules, we examined trafficking of ciM6PR, a cargo that is normally trafficked from endosomes to the trans-Golgi network (TGN) in a retromer complex- and SNX1-tubule dependent manner.[8, 44]

Some ciM6PR is present on the plasma membrane and its endocytosis and trafficking from endosomes to the TGN can be monitored by uptake experiments using antibodies to extracellular epitopes. We therefore incubated our protrudin-depleted and control HeLa cell lines with an appropriate ciM6PR antibody and fixed cells for immunoflourescence microscopy at intervals. In both control and protrudin-depleted cells, ciM6PR was present in a punctate peripheral compartment 10 minutes after uptake (Figure 3A). However, after 30 minutes ciM6PR signal had predominantly moved to a central distribution overlapping with the trans-Golgi marker TGN46 in control cells, whereas in protrudin-depleted cells an increased proportion of ciM6PR remained in puncta (Figure 3A). Consistent with this, after 30 minutes of uptake the percentage of total cellular ciM6PR signal in the TGN46 region of interest was significantly reduced in cells lacking protrudin (Figure 3B). In contrast, total fluorescence of internalised ciM6PR antibody was not significantly different between control and protrudin-depleted cells, indicating that the difference is not due to defective endocytosis of the ciM6PR-antibody complex (Figure 3C). We concluded that defective ETF in cells lacking protrudin impairs endosome-to-Golgi traffic of key receptor cargoes.

**Figure 3.**
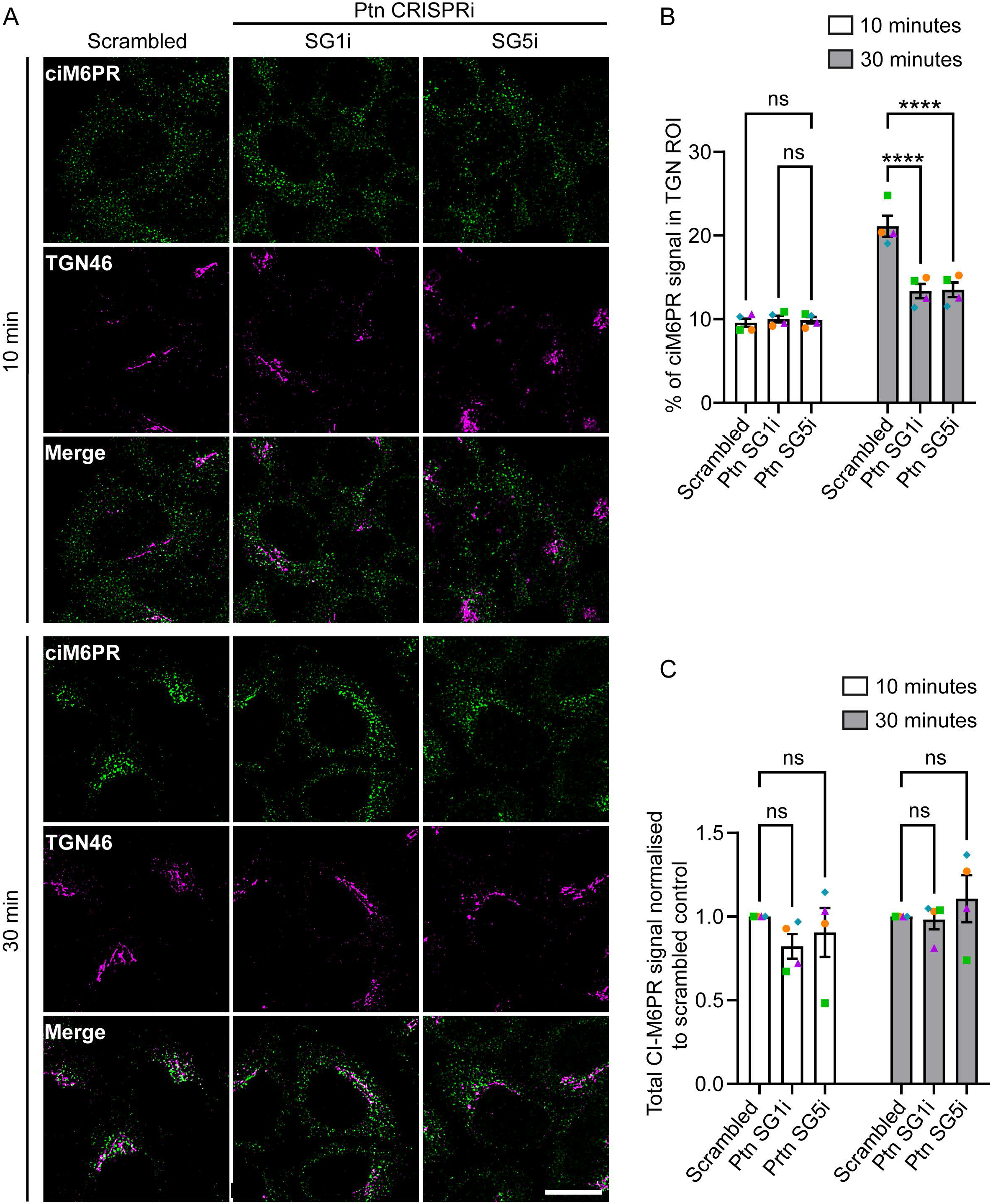
Protrudin depletion delays endosome-to-Golgi trafficking. **A)** The HeLa cell lines indicated were incubated for 10 or 30 minutes with anti-ciM6PR antibody, fixed and visualised for ciM6PR, and endogenous TGN46 to mark the TGN ROI. Scale bar = 20 μm. **B**) The percentage of total cellular ciM6PR in the TGN ROI was quantified. Mean +/- S.E.M. of 4 independent biological repeats are shown; 50 cells per condition per repeat were analysed. **C)** Quantification of total ciM6PR antibody signal per cell for each of the experimental repeats presented in B). For each repeat, signal was normalised to the scrambled control. Means of 4 independent biological repeats +/- S.E.M. are shown; 50 cells per condition per repeat were analysed. Statistical testing was by repeated measures two-way ANOVA with Dunnett correction for multiple comparisons.

### VAP, kinesin and phospholipid-binding domains are required for protrudin to support endosomal tubule fission and endosome-to-Golgi traffic

We next sought to investigate which functional properties of protrudin are required to support ETF. Protrudin has a number domains, motifs and regions that are important for its function and binding to other proteins (Figure 1A). We therefore used retroviral transduction in our CRISPRi-scrambled and CRISPRi-protrudin-SG1i HeLa cell lines to generate stable cell lines expressing myc-tagged wild-type protrudin, myc-tagged forms of protrudin deleted or mutated for key sequence features, or the myc-tag alone (Figure 4A). A low expressing retroviral vector (pLXIN) was used to avoid over-expression artefacts. Proteins deleted for the RAB11 binding region and the low complexity region that binds RAB7 failed to express the relevant protein and were not included in further studies. However, we were able to generate cell lines expressing forms of protrudin deleted for the HDs 1-3 that mediate interactions with other ER shaping proteins (Ptn-ΔHD1-3; residues 67-87, 89-109 and 180-214 deleted), a mutant lacking all N-terminal domains (i.e. the RAB11-binding domain and the HDs; Ptn-Δ1- 208), a FATT motif mutant harbouring an amino acid substitution (D294A) that blocks binding to VAPs (Ptn-FFAT), [39] a mutant lacking the region containing the FFAT and coiled-coil domains that is necessary and sufficient for KIF5 binding (Ptn-ΔFFAT/CC; residues 274-361 deleted),[41] a mutant lacking only the coiled-coil domain that is necessary for KIF5 binding (Ptn-ΔCC; residues 325-345 deleted),[41] and a mutant harbouring two mutations in the FYVE domain (K367A/R369A) that block phosphoinositide binding (Ptn-FYVE) (Figure 4A, B).[45]

**Figure 4.**
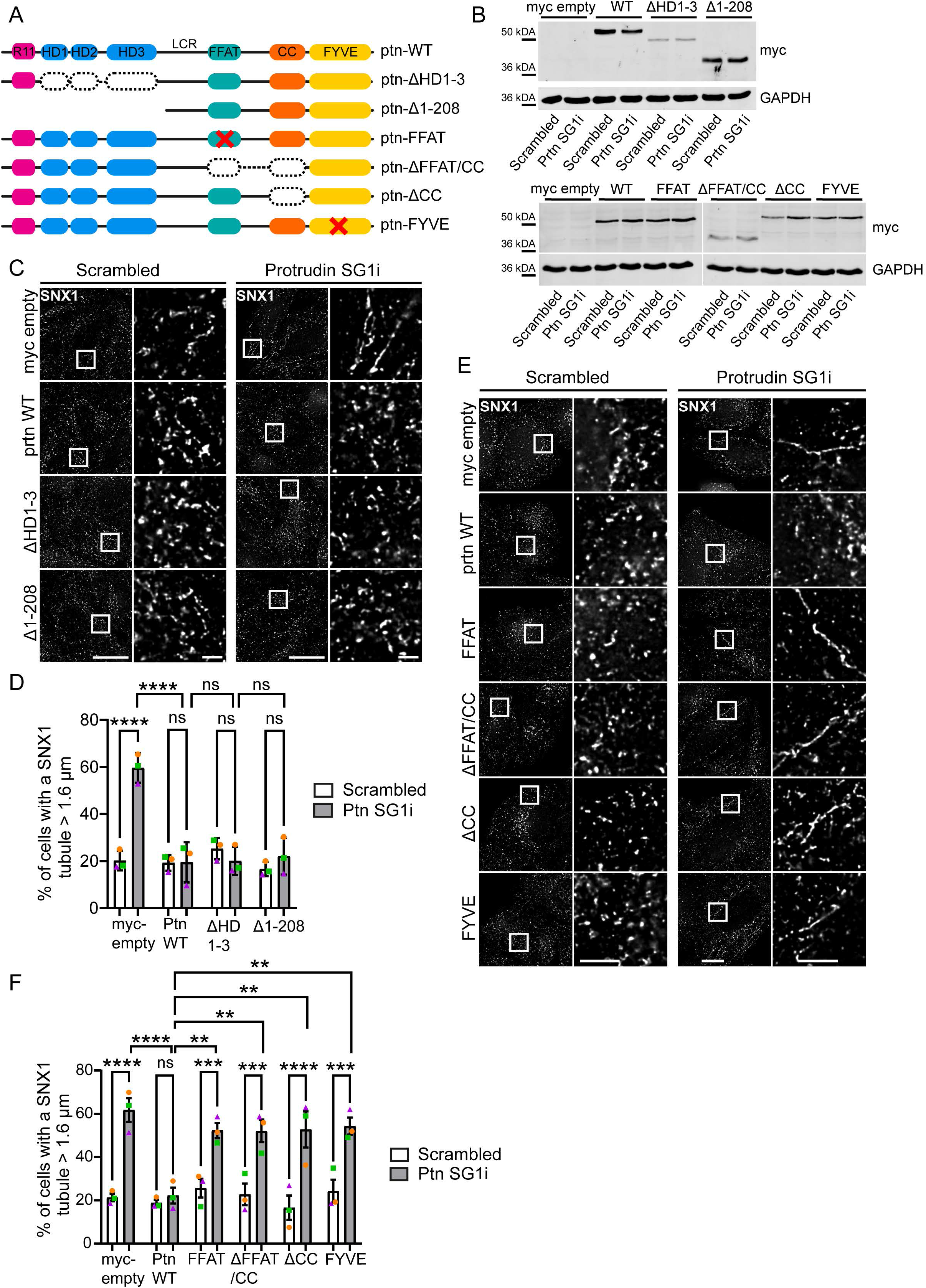
Protrudin requires the ability to interact with VAPs, KIF5 and phospholipids to regulate endosomal tubule fission. **A)** Schematic diagrams showing the mutated versions of protrudin used for the rescue experiments. See Figure 1A for definitions of domain labels. **B**) Immunoblots to validate stable expression of myc-tagged protrudin constructs in scrambled control and protrudin SG1i HeLa cells. **C**) Scrambled control and protrudin SG1i cells stably expressing the protrudin mutant proteins indicated were fixed and labelled for endogenous SNX1. **D**) The percentage of cells with a SNX1-positive tubule longer than 1.6 μm was plotted for each condition. The mean +/- S.E.M. percentage of cells with a long tubule in 3 independent biological repeats is plotted; between 94 and 124 cells per condition per repeat were analysed. **E**) Scrambled control and protrudin SG1i HeLa cells expressing the mutated protrudin proteins indicated were labelled for SNX1. **F**) The percentage of cells with a long SNX1 tubule was quantified in three biological repeats as in D), 100-124 cells per condition per repeat were analysed. In C and E, the inset panel on the right of each column represents a higher magnification view of the boxed area in the corresponding image on the left. Scale bar = 20 μm, inset scale bar = 2 μm. Statistical testing was done by repeated measures two-way ANOVA with Bonferroni correction for multiple comparisons; ns - not significant.

We then used these cell lines to determine the properties of protrudin that are required to rescue the increased endosomal tubulation phenotype seen in cells lacking the endogenous protein. We found that N-terminal domains of the protein were dispensable, as mutants lacking the three HDs (Ptn-ΔHD1-3) or the entire N-terminal region (including the HDs and the RAB11 binding domain; Ptn-Δ1-208) were able to rescue the tubulation phenotype as well as wild-type protrudin (Figure 4C, D). In contrast, all tested C-terminal domains of the protein were critical, as mutants that blocked the ability to bind to VAPs (Ptn-FFAT), to bind to KIF5 (Ptn-ΔCC) and to interact with phosphoinositides all failed to rescue the endosomal tubulation phenotype (Figure 4E, F).

We predicted that similar functional properties would be required for protrudin to support endosome-to-Golgi traffic of ciM6PR, and we tested this with another series of rescue experiments using the ciM6PR uptake assay described above. After 30 minutes of uptake, approximately 20% of total ciM6PR signal is present in the TGN region in scrambled guide control cells expressing the empty myc vector, but this was significantly reduced in cells lacking protrudin (Figure 5A, B). This phenotype was rescued by expression of wild-type protrudin, as well as forms of the protein lacking either the HDs or the entire N-terminal (Figure 5A, B). In contrast, it was not rescued by forms of protrudin that were unable to bind to VAPs, KIF5 or phosphoinositides (Figure 5C, D). In these experiments there was a possible dominant-negative effect of the FYVE mutant protein in the scrambled guide control cells.

**Figure 5.**
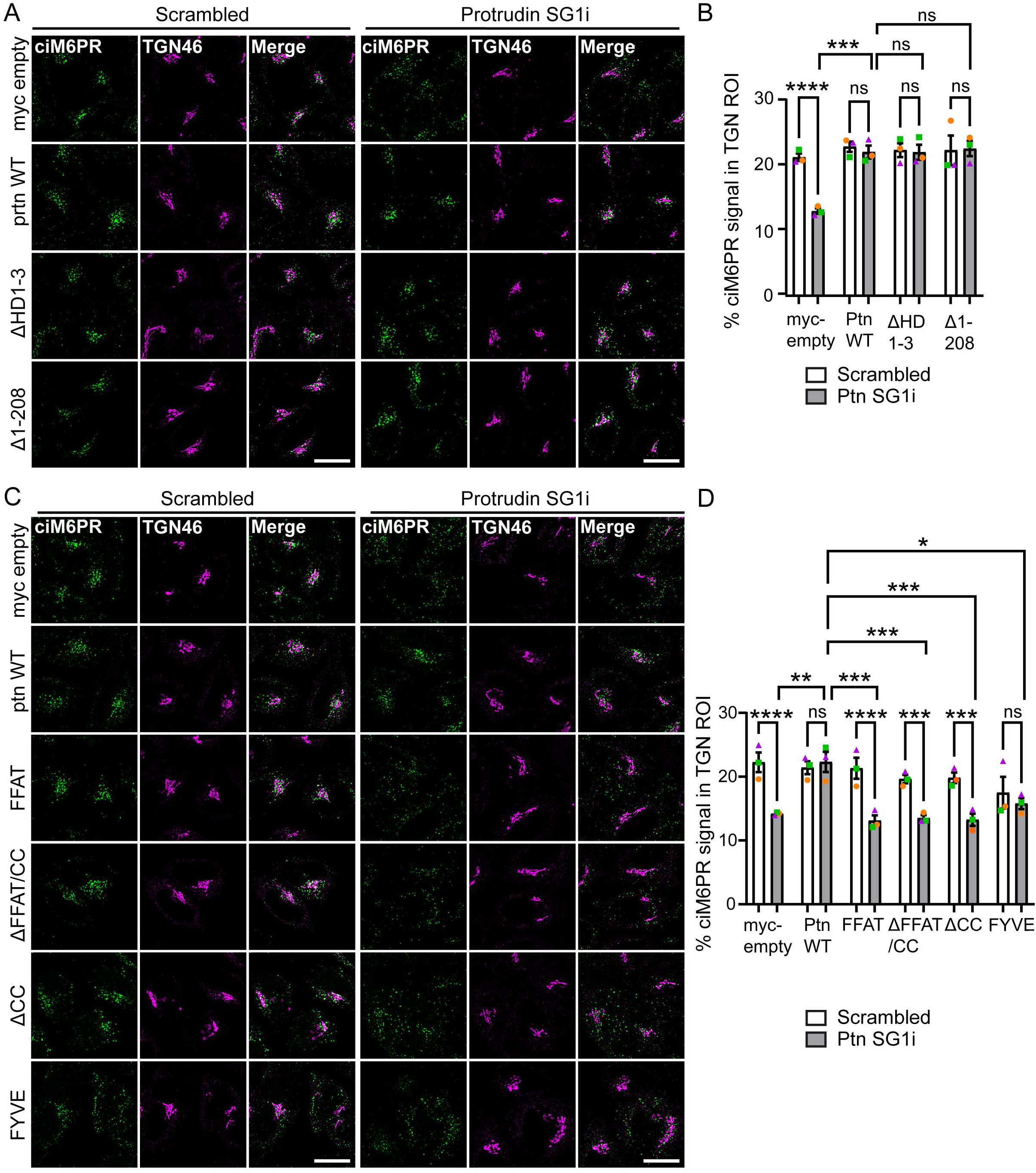
Protrudin requires the ability to interact with VAPs, KIF5 and phospholipids to support endosome-to-Golgi traffic. **A-D)**. Scrambled control and protrudin SG1i cells stably expressing the protrudin mutant proteins indicated were incubated for 30 minutes with anti-ciM6PR antibody, then fixed and visualised for ciM6PR, and endogenous TGN46 to mark the TGN ROI (**A and C**). Scale bar = 20 μm. **B and D**) The mean +/- S.E.M. percentage of total cellular ciM6PR in the TGN ROI 30 minutes after antibody uptake was quantified. N = 3 biological repeats; in B) 56-62 and in D) 49-74 cells per condition per repeat were analysed. Statistical testing was done by repeated measures two-way ANOVA with Bonferroni correction for multiple comparisons.

We concluded that protrudin requires the ability to interact with ER-resident VAPs, endosomal phosphoinositides and KIF5 in order to promote ETF and cargo transport from endosomes to the Golgi apparatus. In contrast, the ability to bind RAB11 was dispensable for these functions of protrudin. As it requires both ER and endosomal interactions, this function of the protein is very likely to take place at an ER-endosome MCS.

### Mutation of key protrudin domains affects the physical distance between protrudin and retromer-positive endosomes

We next sought to better understand the physical relationship between protrudin and SNX1/retromer-positive endosomes. We first analysed whether protrudin co-localises with retromer-related proteins by performing live cell microscopy experiments with stable cell lines expressing mCherry-SNX1 and eGFP-protrudin. We found numerous puncta that showed strong co-localisation between the two markers (Figure 6A, Supplementary Movie 4), and in some cases protrudin signal was seen to surround a SNX1-positive structure (Figure 6B), likely representing ER tubules surrounding an endosome.

**Figure 6.**
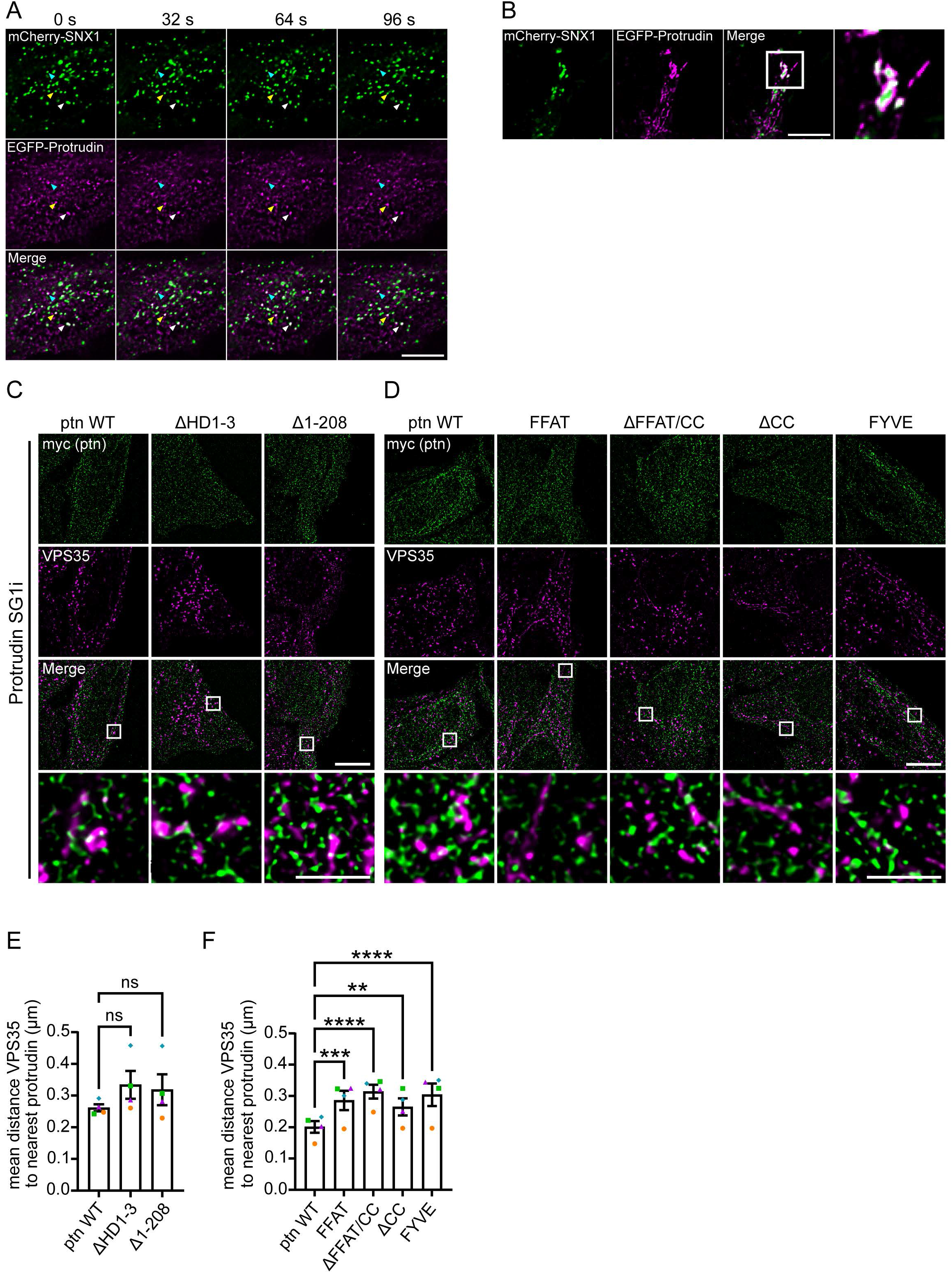
Protrudin co-localises with SNX1 and mutation of protrudin domains affects its distance from retromer-positive endosomes. **A)** Stills from movie (Supplementary Movie 4) showing MRC5 cells stably expressing mCherry-SNX1 and eGFP-protrudin. Arrows mark puncta positive for both mCherry-SNX1 and EGFP-protrudin that traffic together. Movies were acquired using AxioObserver widefield microscope for up to 100 seconds at a frame rate of 500 ms. Scale bar = 10 μm. **B**) Image showing mCherry-SNX1 positive endosomes surrounded by EGFP-protrudin. Scale bar = 10 μm. **C-D**) Representative SIM^2^ images of protrudin SG1i HeLa cells stably expressing the myc-tagged protrudin proteins indicated. The cells were fixed and labelled for endogenous VPS35 and for the myc tag. Scale bar = 10 μm, inset scale bar = 2 μm. **E-F**) Quantification of the mean +/- S.E.M. distance between VPS35- positive endosomes and their nearest protrudin-positive structure. N = 4 biological repeats; 10 cells per condition per repeat analysed. Statistical testing was by repeated measures one- way ANOVA with Dunnett correction for multiple comparisons.

We then explored whether mutation of key protrudin sequences alters the physical distance between protrudin and retromer-positive endosomes. We carried out super-resolution SIM^2^ microscopy between protrudin and the endogenous retromer component VPS35, in stable cell lines expressing myc-tagged protrudin mutants in the background of the CRISPRi- protrudin-SG1i HeLa cell line lacking endogenous protrudin (Figure 6C, D). We then used the DiAna distance analysis plugin for ImageJ to quantify the proximity of each VPS35-structure to the nearest myc-protrudin positive structure. We found that deletion of the HDs or the entire N-terminal half of protrudin, including the HDs and the RAB11-binding region, did not significantly alter the mean distance between VPS35 puncta to the nearest protrudin-positive structure (Figure 6E). In contrast, mutation of the FFAT motif, deletion of the KIF5-binding coiled-coil domain alone or in combination with the FFAT motif, or mutation of the FYVE domain all resulted in a greater mean distance between VPS35 and the nearest protrudin-positive structure (Figure 6F). As all of these mutants retain the HDs that anchor protrudin to the ER, this indicates that these C-terminal motifs are required to regulate the distance at ER-endosome contacts between protrudin-positive ER domains and retromer-positive endosomes. Notably, the properties of protrudin that are required to regulate ER-endosome contact distance are the same as those required to regulate ETF, consistent with the idea that these two functions of protrudin are related.

### Endosomal tubulation in cells lacking protrudin requires dynamic microtubules and dynein and is phenocopied by KIF5 but not FYCO1 depletion

As protrudin required the ability to interact with the microtubule motor KIF5 to promote endosomal fission and endosome-to-Golgi traffic, we next turned our attention to the role of microtubules and microtubule motors in protrudin-dependent ETF.

We first examined the effects of microtubule disassembly or stabilisation. We found that the endosomal tubulation phenotype in cells lacking protrudin was absent in cells in which microtubules had been disassembled by nocodazole (Figure 7A, B) and also in cells where microtubules had been stabilised by taxol (Figure 7C, D). We concluded that the longer ETs seen in cells lacking protrudin require intact and dynamic microtubules for their formation.

**Figure 7.**
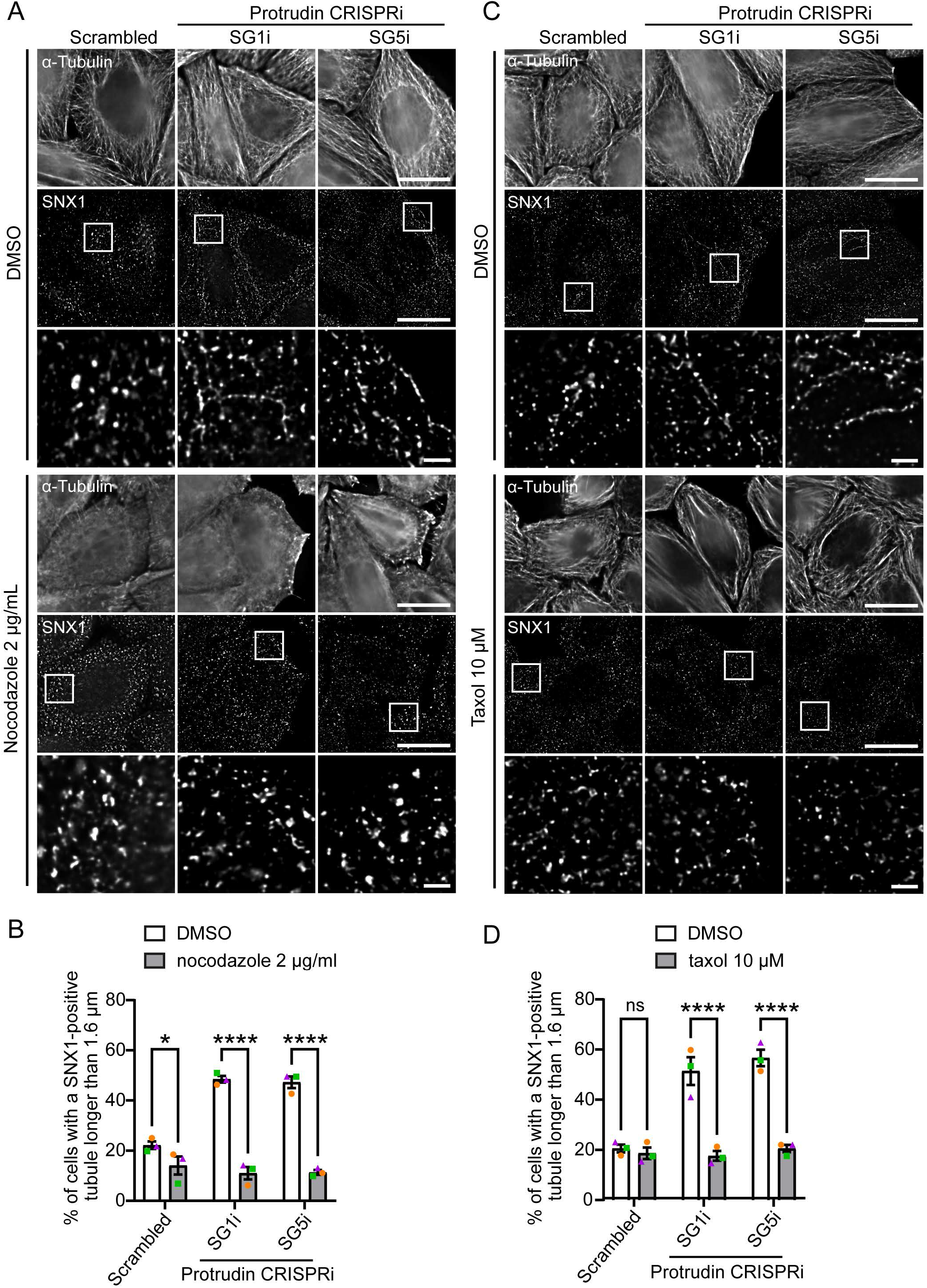
Endosomal tubules induced by protrudin depletion require intact and dynamic microtubules. **A)** The scrambled control and protrudin CRISPRi HeLa lines indicated were treated with 2 μg/mL nocodazole for 60 minutes or with vehicle control (DMSO). Cells were fixed and labelled for endogenous α-tubulin and SNX1. **B**) The mean +/- S.E.M. percentage of cells with a SNX1 tubule > 1.6 μm was plotted. N = 3 biological repeats; 97-119 cells per condition per repeat analysed. **C**) The scrambled control and protrudin CRISPRi HeLa cells indicated were treated with 10 μM taxol for 100 minutes or with vehicle control (DMSO) and labelled as in A). **D**) ETs were quantified as in B), in 3 biological repeats; 100-118 cells per condition per repeat analysed. In A) and C), magnified views of the regions indicated by boxes are shown in the panels below the corresponding image. Scale bars = 20 μm, inset scale bar = 2 μm. Statistical testing in B) and D) was done by repeated measures two-way ANOVA with Bonferroni correction for multiple comparisons.

Next we investigated how KIF5 and the KIF5 adapter protein FYCO1 that is required for protrudin-mediated late endosome motility relate to the role of protrudin in ETF. We hypothesised that protrudin promotes ETF by increasing the KIF5-dependent force acting to break the tubule, and so we predicted that cells lacking KIF5 would also show increased endosomal tubulation indicative of defective tubule fission. We therefore transduced CRISPRi-HeLa cell lines with single guide RNAs targeting KIF5B (the kinesin present in these cells) and selected the two guides with the most efficient depletion for further study (Supplementary Figure 3A). Notably depletion of KIF5B mRNA in these two lines was relatively inefficient (c. 14 % and 36 % of transcript remaining), but even so both lines showed an increase in the proportion of cells that had long ETs (Figure 8A, B), consistent with previous studies that used siRNA-mediated depletion of KIF5.[10] We then tested whether FYCO1 is involved in ETF, using a similar approach to that described above to generate CRISPRi HeLa cell lines depleted of the protein. The two cell lines used had very efficient depletion of FYCO1 at the transcript and protein level (Supplementary Figure 3B, C). However, there was no increase in endosomal tubulation in these cell lines, suggesting that FYCO1 is not required for ETF (Figure 8C, D).

**Figure 8.**
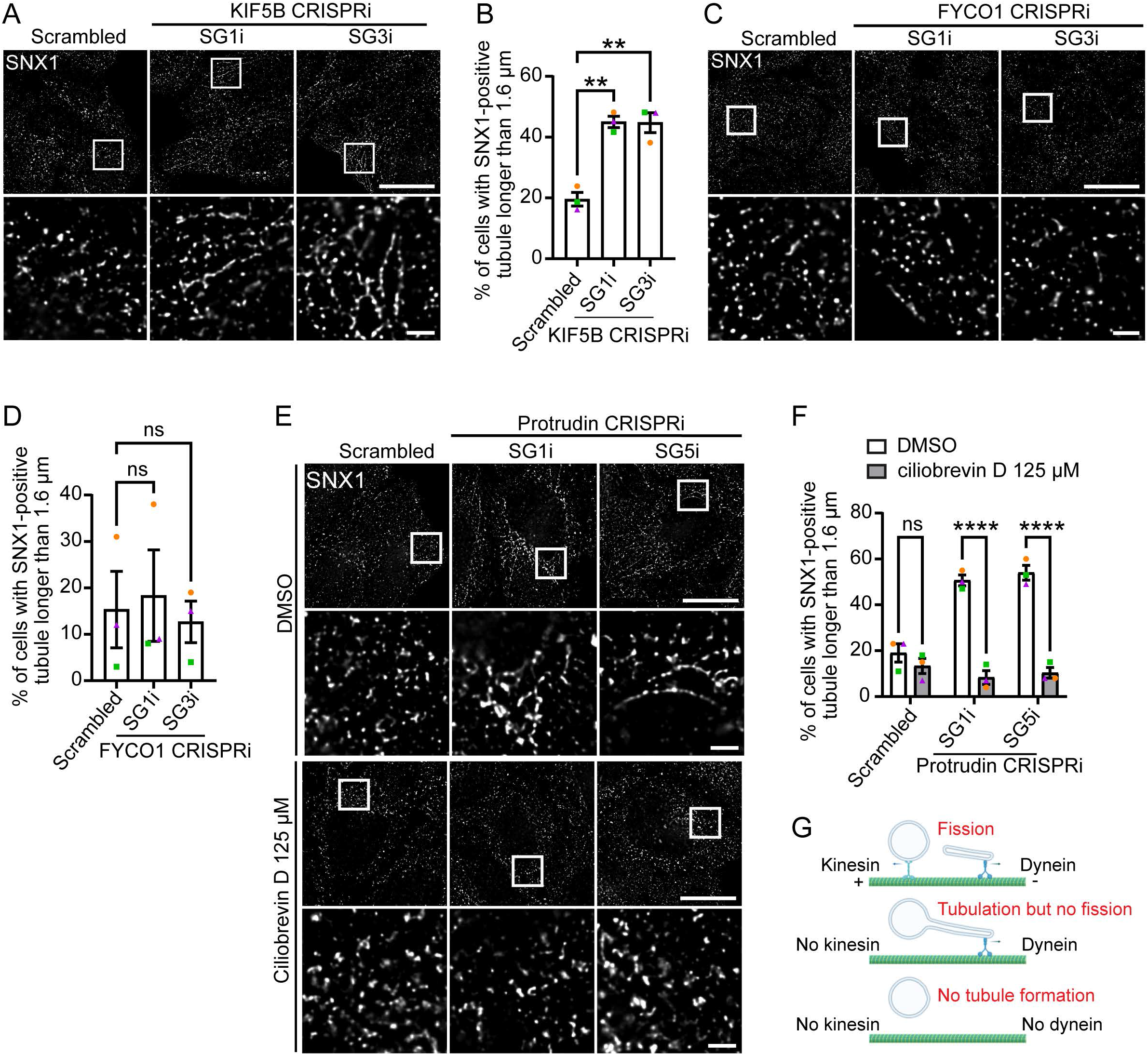
The effect of protrudin depletion on endosomal tubulation is phenocopied by depletion of KIF5 but not FYCO1 and requires dynein. **A)** CRISPRi-HeLa cells stably expressing a scrambled control SG RNA or SG RNAs targeting the KIF5B transcriptional start site (SG1i and SG3i) were fixed and labelled for endogenous SNX1. **B**) The mean +/- S.E.M. percentage of cells with a SNX1-positive tubule > 1.6 μm was plotted. N = 3 biological repeats, 93 - 110 cells per condition per repeat analysed. **C**) Similar immunofluorescence experiments were performed in cells stably expressing SGs targeting the FYCO1 transcriptional start site, and the results are quantified in **D**), which shows the mean +/-S.E.M. percentage of cells with a SNX1 tubule > 1.6 μm for three biological repeats, 100 cells analysed per condition per repeat. **E**) Scrambled control and protrudin CRISPRi HeLa cells were treated with 125 μM ciliobrevin D for 60 minutes or with vehicle control (DMSO), then were fixed and labelled for endogenous SNX1. **F**) The mean +/- S.E.M. percentage of cells with a SNX1 tubule > 1.6 μm was plotted for 3 biological repeats; 100 cells per condition per repeat analysed. **G**) Model for the tug-of-war relationship between KIF5 and dynein- dependent forces in ETF. In the micrographs the lower panel images are magnified views of the boxed areas in the corresponding upper panel image. Scale bar = 20 μm, inset scale bar = 2 μm. Statistical testing was done by repeated measures one-way ANOVA with Dunnett correction for multiple comparisons in B) and D) and by repeated measures two-way ANOVA with Bonferroni correction for multiple comparisons in F).

Elongation and fission of ETs is believed to involve the action of opposing microtubule plus-end (generally kinesin-mediated) and minus-end directed (generally dynein-mediated) motor protein forces. We therefore predicted that, by removing the countervailing force, inhibition of dynein would rescue endosomal tubulation caused by lack of a protrudin/KIF5- generated force. We used the dynein inhibitor ciliobrevin D to test this in cells lacking protrudin and observed complete rescue of the endosomal tubulation phenotype by this drug (Figure 8E, F).[46] TGN dispersal, a known consequence of dynein inhibition, was used as a positive control for the effects of the drug on dynein (Supplementary Figure 3D).

We concluded from these experiments that protrudin likely recruits KIF5 to provide a microtubule-dependent force that promotes ETF, in concert with a counterbalancing dynein-dependent force (Figure 8G). This process does not involved FYCO1, and so is mechanistically distinct from protrudin’s role in late endosome transport.

### Protrudin is developmentally regulated and controls endosomal tubulation in human cortical neurons

Protrudin promotes neurite outgrowth and axonal regeneration and so we examined whether protrudin’s role in regulating ETF was relevant to human neurons. To do so we used the i^3^ (integrated, inducible and isogenic) neuron system.[47] In this system the NGN2 neurogenic transcription factor gene, under a doxycycline responsive promotor, is knocked-in at a safe harbour locus in the well-characterised stem cell line WTC11. Incubation with doxycycline allows rapid, scalable and reproducible differentiation of the i^3^ iPSCs to glutamatergic cortical neurons (i^3^Ns) within 14 days. We first characterised protrudin expression during a time-course of iPSC differentiation to neurons. At the RNA and protein level, upregulation of protrudin expression began at day three after initiation of differentiation and steadily increased up to day 21 (Figure 9A-B, Supplementary Figure 4A, B), indicating that it is developmentally regulated.

**Figure 9.**
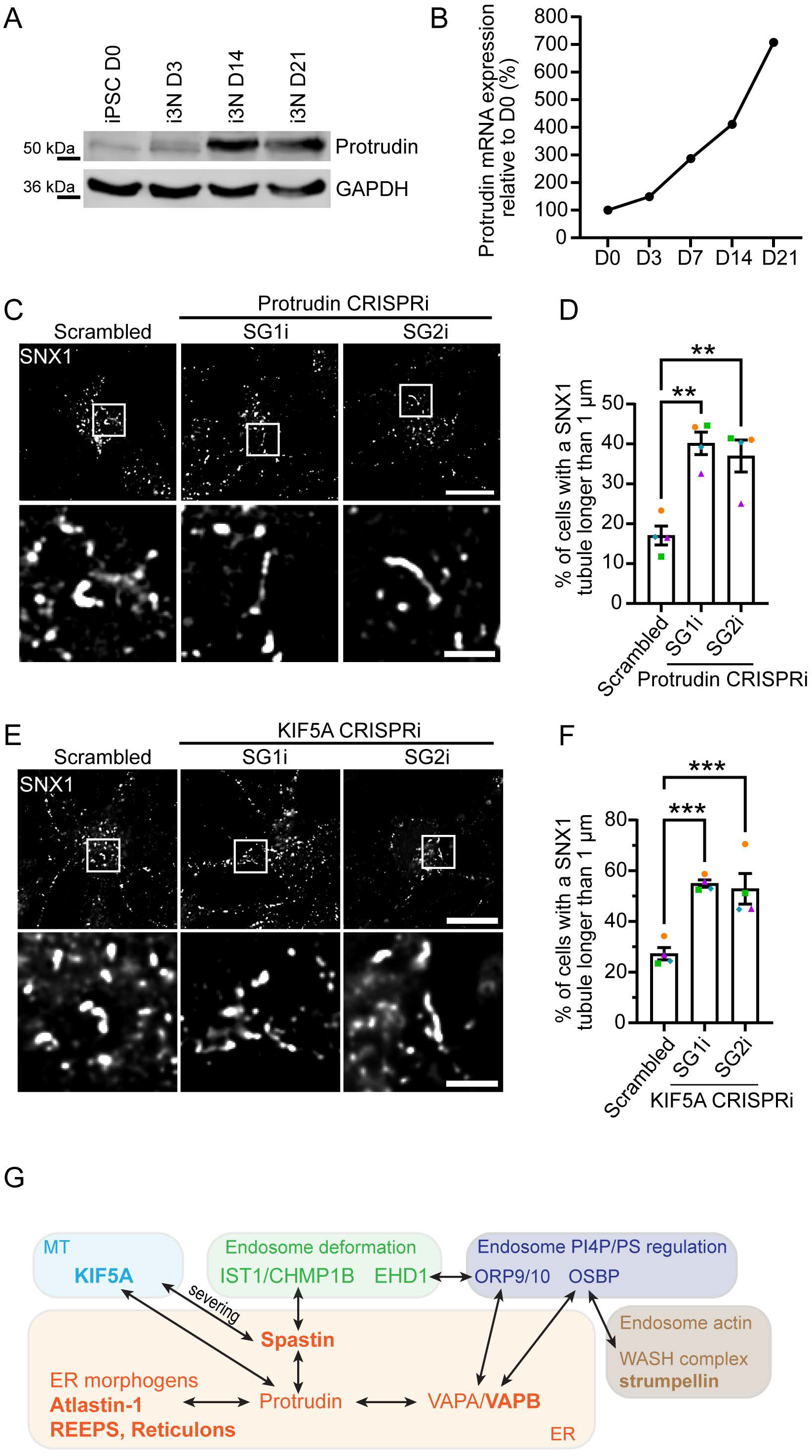
Protrudin is developmentally regulated and protrudin or KIF5 depletion causes increased endosomal tubulation in human cortical neurons. **A)** I^3^ iPSCs were differentiated to I^3^Ns over a 21 day time course. Cell lysates were prepared 3 (D3), 14 (D14) or 21 (D21) days after differentiation was induced and then immunoblotted for protrudin. GAPDH blotting serves to validate equal lane loading. **B)** RT-qPCR quantification of protrudin mRNA expression in I^3^Ns over a differentiation time-course. *ACTB* was used as a housekeeping control to normalise results; protrudin expression was normalised to Day 0. N = 1 biological repeat, each consisting of three technical replicates. **C)** Scrambled control and protrudin CRISPRi day 14 I^3^Ns were fixed and labelled for endogenous SNX1. **D)** The mean +/- S.E.M. percentage of cells with a SNX1 tubule > 1 μm was plotted for 4 biological repeats; 51 - 103 cells per condition per repeat analysed. **E)** Scrambled control and KIF5A CRISPRi day 14 I^3^Ns were fixed and labelled for endogenous SNX1. **F)** The mean +/- S.E.M. percentage of cells with a SNX1 tubule > 1 μm was plotted for 4 biological repeats; 68 - 91 cells per condition per repeat analysed. **G)** Schematic diagram of processes involved in ETF that are linked by protrudin. In micrographs the scale bar = 10 μm, inset scale bar = 2 μm. Statistical testing in D) and F) was done by repeated measures one-way ANOVA with Dunnett correction for multiple comparisons.

To generate human neurons lacking protrudin we used a version of the i^3^ iPSC line that is integrated for CRISPRi machinery at a second safe harbour locus.[48] Using lentiviral transduction of sgRNAs targeting the *ZFYVE27* transcriptional start site we generated two i^3^ iPSC lines lacking protrudin (SG1i and SG2i) as well as a control line transduced with a scrambled sgRNA. We confirmed >98% protrudin depletion in these cell lines at the mRNA level (Supplementary Figure 4C)). We then differentiated these lines to day 14 neurons, confirmed that protrudin depletion was maintained (Supplementary Figure 4D) and examined endosomal tubulation of SNX1 by immunofluorescence microscopy. We found a significant increase in the percentage of cells with long ETs (here defined as tubules longer than 1 μm, as ETs are smaller in these cells than in HeLa or MRC5 cells) in both cell lines depleted of protrudin (Figure 9C, D).

Finally, in view of our results with KIF5B, we examined whether depletion of KIF5A, the main form of KIF5 found in central nervous system neurons, affected endosomal tubulation. Using the CRISPRi-i^3^ iPSC line, we generated two lines lacking KIF5A (Supplementary Figure 4E, F) and differentiated them to 14 day old neurons. Again we found that both KIF5A-depleted cell lines had an increase in the percentage of cells with long ETs (Figure 9E, F).

We concluded that protrudin and KIF5 are required for the correct regulation of endosomal tubulation in human neurons. In the context of the work above this very likely represents a role for these proteins in neuronal ETF.

## Discussion

In this study we show that protrudin is required for efficient fission of endosomal transport tubules and onwards traffic of receptors, such as ciM6PR, that are normally transported on them. This function of protrudin almost certainly happens at ER-sorting endosome contacts, as, a) it required the capacity of protrudin to interact with both ER-localised VAP proteins via the FFAT motif and endosomal membrane phospholipids via the FYVE domain, b) full-length protrudin, which is known to localise to the ER, and the ET marker protein SNX1 showed strong colocalisation, and c) point mutations that render the FFAT motif and FYVE domain non-functional increased the mean distance between protrudin and the retromer component VPS35, as would be expected after disruption of an ER-endosome contact. A role for protrudin at ER-sorting endosome contacts is consistent with the previous observation that protrudin also participates in contact sites between ER and late endosomes, in a process that governs late endosomal motility and position.[40]

ETF involves a tug-of-war between opposing kinesin- and dynein- dependent forces, which is believed to provide mechanical force to help break the ET.[10, 11] Protrudin required the ability to bind to KIF5 to promote ETF, while the endosomal tubulation phenotype observed on protrudin depletion was phenocopied by KIF5 depletion. These data are consistent with the idea that protrudin promotes KIF5 loading onto SNX1-positive endosomes, analogous to its role in late endosomal motility. Failure of ETF in cells lacking protrudin blocked ciM6PR traffic out of endosomes. This illustrates that in the tug-of-war model, deficient microtubule + end-directed transport (e.g. caused by lack of protrudin or KIF5) can also result in defective – end-directed transport of cargoes, such as ciM6PR, that normally traffic in a dynein-dependent manner, if the ETs that carry them are rendered unable to fission from the parent endosome.[49] In late endosome motility, protrudin is proposed to hand over KIF5 to the endosomal kinesin adaptor FYCO1, and this protein is also required for late endosome motility.[40] However, FYCO1 depletion did not cause an increase in endosomal tubulation, indicating that it is not involved in ETF. Further work will be required to understand whether another endosomal KIF5 adapter is involved in the ETF process, or whether KIF5 binds directly to the endosome. Nevertheless, these data suggest that protrudin participates in at least two spatially (i.e. early sorting endosomes and late endosomes) and mechanistically distinct KIF5-dependent processes in endosomal trafficking and transport.

The N-terminal RAB11 binding domain was dispensable for protrudin’s role in ETF. This region is proposed to interact preferentially with inactive, GDP-bound RAB11 and retain it in this state, so switching off generalised cellular recycling through the RAB11-positive perinuclear recycling endosomal compartment.[37] These endosomes are physically distinct from the early sorting endosome and this likely explains why RAB11 interaction is not required for ETF from sorting endosomes. Perhaps more surprising was our finding that the HDs of protrudin, which serve to anchor it in the ER membrane, are dispensable for ETF. However, it should be noted that the FFAT motif, which interacts with ER-localised VAP proteins, was required for ETF, and so the capacity to interact with the ER is important for this function of protrudin. Furthermore, our rescue experiments did involve modest over- expression of protrudin constructs, and so it may be that in the physiological situation the HDs do play some role in guiding protrudin to localise to ER tubular domains, so facilitating the interaction with VAP proteins that is critical for ETF.

This work adds another piece to our understanding of the mechanisms of ETF. A picture is emerging in which protrudin is a central hub of this process (Figure 9G). Protrudin binds to VAPS, and interestingly protrudin is the only ER-resident membrane protein known to have a FFAT motif, making it uniquely placed to recruit VAP proteins to the highly curved ER tubules where ETF takes place.[50] VAPs interact with OSBP and ORP9/10, which respectively regulate endosomal PI4P and phosphatidylserine concentrations, thereby mediating WASH complex recruitment to generate actin on the endosome and insertion of the scission protein EHD1.[23, 24] Protrudin also interacts with KIF5 and with spastin, whose microtubule severing activity is required for ETF, possibly by facilitating dynein loading at new microtubule plus ends.[22, 51] Thus protrudin can co-ordinate proteins on both sides of the motor protein “tug of war” that is involved in ETF. By interacting both with these microtubule-associated proteins and with VAPS, protrudin links the lipid-modifying, actin-modifying and microtubule-associated machineries involved in ETF. Furthermore, through a largely unknown mechanism, correct ER morphogenesis is also required for ETF.[21, 32] Protrudin interacts with ER morphogens, including atlastin-1 and members of the reticulon and REEP families, thereby linking this process to the other mechanisms involved in ETF and perhaps suggesting that abnormality of these morphogens could affect ETF by disrupting protrudin’s correct functioning.[36] Finally, the interaction between spastin and IST1 also serves to link ET membrane deformation by the IST1-CHMP1B atypical ESCRT complex to these other processes involved in ETF, via spastin’s interaction with protrudin. Thus, protrudin is a hub for processes involved in ETF.

ETF is highly pathologically relevant, as multiple genes that encode proteins involved in this process are mutated in hereditary motor neurons disorders. While an alteration in the protrudin gene was initially thought to be a cause of HSP, it is now clear from the Gnomad database that the variant described is a common general population polymorphism with an allele frequency of approximately 1 in 100.[52] However, mutations in the genes encoding spastin, atlastin-1, the WASH complex member WASHC5 and KIF5A cause autosomal dominant HSP, while mutations affecting the membrane scission protein dynamin 2 cause hereditary motor and sensory neuropathy and HSP, and mutations in VAPB cause motor neuron disease.[29, 30, 33, 34, 53–55] Thus the machinery of ETF is highly enriched for proteins that are implicated in hereditary motor neuron disorders. Signs of defective ETF have been found in human or animal neuronal models of spastin and atlastin-1 HSP,[12, 22, 32] and in this study we show that lack of KIF5A also causes increased endosomal tubulation in human neurons.

How is the role of protrudin in ETF related to its other functions in promoting neurite extension/protrusion formation and axonal regeneration? Mutation of the FFAT motif or FYVE domain, or removal of the KIF5-binding coiled-coil domain along with the FFAT motif, all reduced protrudin’s ability to promote axonal regeneration, while proper function of these domains is required for protrusion formation in non-polarised cells and promotion of neurite growth.[37–41, 45, 56] Thus the domains and functional properties that were required for protrudin to promote ETF are also required for these other roles of protrudin, suggesting that ETF could contribute to them. A possible mechanism for this could be, for example, by promoting the axonal trafficking of nerve growth molecules such as NgCAM, which are endocytosed in the somatodendritic region and transcytosed to the axon, via a pathway that involves ETs.[57] Interestingly, the intracellular distribution of NgCAM is known to be altered in cells lacking protrudin.[37] However, properties of protrudin that are not required for ETF are also important for its other functions. The RAB11 binding domain is required for protrusion formation and neurite extension, while the HDs are required for these processes and also axonal regeneration.[37–39] Thus, ETF may contribute to these functions of protrudin, but they must also require other pathways, such as altered RAB11- dependent endosomal traffic or protrudin-promoted late endosomal transport via FYCO1/KIF5 in promoting protrusion formation, or protrudin-promoted accumulation of ER and RAB11 endosomes in the growth cone in axonal regeneration.

In summary, in this study we demonstrate that protrudin directly promotes ETF via a KIF5- dependent mechanism at ER-endosome contacts. Through its many interactions protrudin can act as a key hub to co-ordinate many other molecular machineries that are involved in ETF.

## Methods

### Antibodies

Primary antibodies used for western blotting (WB) were: in-house rabbit polyclonal anti-spastin (86-340, raised against a glutathione S-transferase fusion protein that incorporates residues 86-340 of M1-spastin) produced as previously described,[58] rabbit polyclonal anti- ZFYVE27 (Proteintech, 12680-1-AP), rabbit polyclonal anti-Sox2 (Cell Signalling Technology, 2748), rabbit polyclonal anti-Oct4 (Cell Signalling Technology, 2750), mouse monoclonal anti-Nanog [1E6C4] (Cell Signalling Technology, 4893), mouse monoclonal anti-Tau (Abcam, ab80579), mouse monoclonal anti-MAP2 [AP-20] (Abcam, ab11268), rabbit polyclonal anti-Beta-III tubulin (Abcam, ab18207), rabbit polyclonal anti-GFP (Abcam, ab6556), mouse monoclonal anti-SNX1 (BD Transduction Laboratories, 611482), mouse monoclonal anti-myc tag [4A6] (EMD Millipore, 05-724), mouse monoclonal anti-M6PR (cation independent) [2G11] (Abcam, ab2733), rabbit monoclonal anti-GAPDH [14C10] (Cell Signalling Technology, 2118). IRDye-conjugated secondary antibodies for WB were purchased from LICOR.

Primary antibodies for immunofluorescence microscopy were: mouse monoclonal anti-alpha tubulin (acetyl K40) [6-11B-1] (Abcam, ab24610), mouse monoclonal anti-SNX1 (BD Transduction Laboratories, 611482), mouse monoclonal anti-M6PR (cation independent) [2G11] (Abcam, ab2733), Alexa Fluor 647-conjugated mouse monoclonal anti-M6PR (cation independent) [2G11] (Abcam, ab205813), mouse monoclonal anti-myc tag [4A6] (EMD Millipore, 05-724), goat polyclonal anti-VPS35 (Abcam, ab10099), sheep polyclonal anti- TGN46 (Bio-Rad, AHP500G). Alexa Fluor-conjugated secondary antibodies for IF were obtained from Molecular Probes.

### Constructs

For CRISPRi, sgRNA sequences targeting the transcription start site of protrudin, FYCO1, KIF5A, and KIF5B were selected from the Weissman CRISPRi-v2 library or designed for this study using the CRISPick online tool.[59–61] SgRNA sequences used were: Protrudin SG1i GTGCGACTTCCGAACAACCC; Protrudin SG2i GAGGCCGAGCCAAGGCGAAA; Protrudin SG5i GAGGAGCGGCGTTCAGCCGG; FYCO1 SG1i GGGGCCAAGCGGAAGAGGAC; FYCO1 SG3i GGACGGGAACGCGGTGCTGA; KIF5A SG1i GGCGCGCTGTCTCTCTCCTG; KIF5A SG2i GCCGAAAGGACCAGACGCCC; KIF5B SG1i GGGCGGCCTCAGGAGTGATC; KIF5B SG3i GTCCCTGCAAGACTGAGCGG; Scrambled GGGACGCGAAAGAAACCAGT.

Sense and antisense sgRNA oligonucleotides were designed with 5’CACC and 3’CAAA overhangs and cloned into pKLV U6gRNA-EF(BbsI)-PGKpuro2ABFP digested with BbsI or lenti-sgRNA hygro digested with BsmBI. pKLV U6gRNA-EF(BbsI)-PGKpuro2ABFP was a gift from Kosuke Yusa (Addgene plasmid #50946) and lenti-sgRNA-hygro was a gift from Brett Stringer (Addgene plasmid #104991). Lenti-dCas9-KRAB-blast was a gift from Gary Hon (Addgene plasmid #89567).

Lentiviral vectors for protrudin and SNX1 expression used the pLV-EF1a-IRES-Hygro (gift from Tobias Meyer, Addgene plasmid #85134) and the A62 backbone (gift from Michael Fernandopulle, University of Cambridge), respectively. EGFP-protrudin (gift from James W. Fawcett, University of Cambridge) was cloned into pLV-EF1a-IRES-Hygro using Gibson assembly and mCherry-SNX1 (gift from Peter J Cullen, University of Bristol) was cloned into the A62 vector also using Gibson assembly.

Retroviral plasmids for protrudin expression in HeLa cells used the pLXIN backbone (gift from Andrew Peden, University of Sheffield). pLXIN-myc-protrudin constructs were generated by cloning protrudin (*ZFYVE27,* UniProt reference Q5T4F4-3) and its mutated and truncated versions from AAV-syn-EGFP-ZFYVE27 (gift from James W. Fawcett, University of Cambridge) into the pLXIN vector (EcoRI-BamHI) using Gibson assembly with the addition of an N-terminal myc tag.[38] pLXIN-myc-protrudin Δ1-208 was created specifically for this study by truncating the WT protrudin sequence using PCR. A control construct expressing myc tag alone was created by cloning annealed oligos with the myc tag sequence into the pLXIN vector. pMD VSV-G and pCMV Δ8.91 envelope and packaging vectors were used for lentiviral transductions and pMD.GagPol and pMD.VSVG for retroviral transductions.

### Cell culture

HeLa, MRC-5, and HEK 293T cells were cultured at 37 °C and 5 % CO2 in complete Dulbecco’s Modified Eagle Medium with 4500 mg/L glucose, sodium pyruvate, and sodium bicarbonate (DMEM, Sigma-Aldrich D6546) supplemented with 10 % fetal bovine serum (FBS, Sigma-Aldrich, F7524), 2 mM L-glutamine (Sigma-Aldrich, G7513), 100 U/mL penicillin and 100 µg/mL streptomycin (Sigma-Aldrich, P0781). If cells were to be transfected, transduced, or selected with other antibiotics, penicllin and streptomycin were omitted from the media. Cells were passaged using Trypsin-EDTA (Sigma-Aldrich, T3924). Cells were regularly tested for mycoplasma contamination using EZ-PCRTM Mycoplasma Detection Kit (Geneflow, K1-0210).

Human induced pluripotent stem cells (iPSCs) were cultured at 37 °C and 5 % CO2 in Essential 8 medium (E8, ThermoFisher Scientific, A1517001) on plates coated with Matrigel (Corning, 354277). E8 medium was replaced daily and iPSCs were passaged at 80 % confluence with 0.5 mM EDTA. StemBeads FGF2 (StemCultures, SB500) were added to E8 media occasionally to allow media change-free culture for up to 72 hours. RevitaCell Supplement (Gibco, A2644501) was added to iPSCs after thawing to improve recovery and ROCK inhibitor Y-27632 (10 µM, Tocris, 1254) was added occasionally after passaging to improve viability. Cells were cryopreserved in KnockOut Serum Replacement. All iPSC lines were tested regularly for mycoplasma contamination EZ-PCRTM Mycoplasma Detection Kit.

### Generation of Stable cell lines

#### HeLa cells stably depleted of spastin, protrudin, FYCO1, and KIF5B

These were generated via sequential lentiviral delivery of CRISPRi components. First, cells were transduced with dCas9-KRAB. Lentiviral particles were generated by co-transfection of HEK 293T packaging cells with a lentiviral expression construct containing dCas9-KRAB and packaging constructs pMD VSV-G and pCMV Δ8.91 at a ratio of 1:0.7:0.3 using TransIT-293 transfection reagent (Mirus Bio, 2704). Media containing the lentivirus were collected 48 h post-transfection, passed through a 0.45 µm filter, and added to target HeLa cells with 10 µg/mL polybrene (Sigma-Aldrich, TR-1003). Successfully transduced cells were selected by adding blasticidin 2.5 µg/mL to the cell culture media for at least 48 hours and dCas9 expression was verified by immunoblotting. HeLa cells stably expressing dCas9-KRAB were then transduced for a second time using the protocol described above, this time with a lentiviral vector containing appropriate sgRNA. Successfully transduced cells were selected by adding puromycin 2 µg/mL for at least 48 hours or hygromycin 200 µg/mL for at least 72 hours. Silencing of target genes was validated at mRNA level using RT-qPCR and at protein level using immunoblotting.

#### Protrudin rescue cell lines

HeLa cells depleted of protrudin with CRISPRi were transduced with retrovirus containing either wild-type protrudin, various mutated and truncated versions of protrudin, or myc tag alone. Retroviral constructs for protein expression used the pLXIN backbone. HEK 293T cells were transfected with pLXIN, packaging and envelope plasmids using TransIT-293 transfection reagent (Mirus Bio, 2704), in 1:0.7:0.3 ratio.

Retrovirus-containing media was harvested 24 hours later and passed through a 0.45 µm filter. Target cells were transduced with retrovirus-containing media diluted 1:1 with fresh DMEM with addition of 10 µg/mL polybrene.

#### GFP-SNX1 cells

MRC-5 cells stably expressing GFP-SNX1 were generated as described previously.[22] The cells were then lentivirally transduced with dCas9-KRAB and sgRNAs (scrambled or targeting protrudin) as described above. MRC-5 cells stably expressing EGFP- protrudin and mCherry-SNX1 were generated by lentiviral transduction, as described above, and selected with hygromycin and puromycin.

#### I^3^ iPSCs stably depleted of protrudin and KIF5A

WTC11 iPSCs with integrated dCas9-BFP-KRAB (gift from Michael Ward, NIH) were transduced with lentivirus containing sgRNA targeting transcription start sites of protrudin or KIF5A, as described above. Briefly, HEK- 293T cells were co-transfected with a lentiviral expression construct and the packaging vectors pCMVΔ8.91 and pMD VSV-G at a ratio of 1:0.7:0.3 using TransIT-293 (Mirus Bio) as per the manufacturer’s instructions. The viral supernatant was collected 48 h post-transfection, passed through a 0.45 µm filter, and added to target cells in the presence of 10 µg/mL polybrene (Sigma-Aldrich). Typically, following spinoculation at 1800 rpm for 1h at 32°C, cells were transduced for 16 h. Transduced cells were selected by adding puromycin at a final concentration of 1 µg/mL from 24 h if required.

### Immunofluorescence microscopy on fixed cells

Cells were grown on glass coverslips to approximately 60% confluence and fixed with 4% (v/v) formaldehyde in PBS for 20 minutes. Cells were then permeabilised with 0.1% (v/v) saponin in PBS for 30 mins and blocked with 3% (w/v) bovine serum albumin with 0.05% (v/v) saponin for 1 hour. Cells were incubated with primary and secondary antibodies for 1 hour each and then incubated 1 µg/mL DAPI for 4 minutes and mounted onto glass slides using Prolong Gold antifade agent (Molecular Probes). Slides were left to cure overnight at room temperature.

Cells were visualised using Zeiss AxioImager Z2 Motorized Upright Microscope or LSM780 confocal microscope and if necessary (e.g. for SNX1 tubule visualisation) were deconvolved before analysis.

### Live-cell microscopy

Cells were grown to 60% confluence in 35 mm glass-bottom dishes (MatTek, P35G-1.5-14-C) and imaged live in Live Cell Imaging Solution (Invitrogen, A14291DJ) on an AxioObserver Z1 inverted microscope with 37°C incubation. Imaging was performed with 500 ms exposure time per frame continuously for 85-90 seconds. Movies were deconvolved before analysis. For quantification of tubule maximum duration 5 tubules per cell in 5 cells (25 tubules total) were analysed per condition per experimental repeat, blind to the experimental condition. Movies were made with Adobe After Effects 2024 and Adobe Media Encoder 2024 software and rendered at 30 frames per second.

### Immunoblotting

Cells were washed on ice with PBS and scraped with ice-cold lysis buffer (1% Triton X-100, 300 mM NaCl, 50 mM Tris [pH 8.0], 5 mM EDTA [pH 8.0], and protease inhibitors), left on ice for 15 min and centrifuged at 10,000 x g for 5 min at 4°C. Sample buffer was added to the supernatant and samples were boiled at 98°C for 5 min. Proteins were resolved by SDS-PAGE and transferred to a PVDF membrane. Membranes were blocked with 5% (w/v) skimmed milk powder in TBS-T (Tris buffered saline 1x with 0.1% Tween-20) with 0.1% Tween-20 for 1 hr at room temperature and incubated with primary antibody overnight at 4°C. Subsequently, membranes were washed with TBS-T 3x 10 min and incubated with an appropriate secondary antibody for 1 hr at room temperature covered from light. Membranes were washed again with TBS-T and imaged directly on an Odyssey Infrared Imaging System using LICOR Image Studio software.

### RT-qPCR

Cells grown in a monolayer were homogenised using QIAshredder (Qiagen) and RNA was isolated using RNeasy kit (Qiagen). cDNA was prepared using high capacity RNA-to-cDNA kit (Thermo Fischer Scientific). Real time quantitative PCR (RT-qPCR) reactions were carried out using 5 ng of cDNA, PCR Master Mix (Thermo Fischer Scientific) and TaqMan Gene Expression

Assays (FAM) (Thermo Fischer Scientific) specific for spastin (*SPAST*), protrudin (*ZFYVE27*) and beta-actin (*ACTB*). Relative gene expression was calculated using the delta-delta Cq method normalised to *ACTB* as a housekeeping gene.

### ciM6PR uptake assay

HeLa cells were plated on glass coverslips and grown to ∼60% confluence. Cells were incubated with an antibody targeting extracellular epitopes of ciM6PR (ab2733) diluted 1:200 in serum-free DMEM for 10 or 30 minutes, washed 3 times with PBS to remove unbound antibody, and fixed with 4% formaldehyde for 20 minutes. Cells were then stained with anti- TGN46 antibody (1:100) and appropriate secondary AlexaFluor antibodies (1:300) to detect ciM6PR and TGN46 primary antibodies. Cells were imaged on a Zeiss LSM880 confocal microscope. To measure ciM6PR distribution, Golgi region of interest (ROI) was selected manually based on TGN46 distribution and ciM6PR signal within that ROI was measured to give Golgi ciM6PR signal. All signal within the cell outside of the Golgi ROI was defined as vesicular ciM6PR and the ratio of vesicular/Golgi ciM6PR was calculated.

### Image analysis

If required (e.g. for SNX1 tubule analysis), images of fixed cells were deconvolved using Huygens Professional version 18.04 (Scientific Volume Imaging). Image analysis was carried out using Fiji (ImageJ). Images were randomised for blind analysis using Blind Analysis Tools plugin in Fiji. Distance and proximity analysis on super-resolution images was conducted using DiAna plugin for ImageJ.[63] A particle size threshold of 50 pixels was set to exclude background noise and nearest neighbour distance analysis was carried out. Figures were prepared in Adobe Photoshop and Illustrator.

### I^3^ neuron differentiation

Differentiation of i^3^ neurons was performed as described by [62]. iPSC WTC11 cells stably expressing neurogenin-2 under a tetracycline inducible promoter were cultured in E8 medium on Matrigel-coated plates.[48] At the start of the differentiation protocol iPSCs were single-cell dissociated with StemPro Accutase (ThermoFisher Scientific, A1110501) and seeded at a density of 1.5 x 10^5^ cells/cm^2^ on Matrigel-coated plates in Induction Medium (IM) with addition of doxycycline hyclate in PBS (2 mg/mL, Sigma-Aldrich, D9891) and ROCK inhibitor Y-27632 (only on day 0 of differentiation). Pre-differentiated cells were cultured for 3 days with daily IM changes. On day 3 of the differentiation protocol, cells were dissociated with StemPro Accutase and replated onto plates coated with 0.1 mg/mL poly-L-ornithine hydrobromide (Sigma-Aldrich, P3655) at a density ranging from 5 x 10^4^ cells/cm^2^ to 5 x 10^5^ cells/cm^2^ depending on application. Following replating, neurons were maintained in Cortical Neuron Culture Medium (CM) for up to 14 days with half-volume CM change every 3-4 days.

## Data analysis and statistics

Statistical analyses were done by one- or two- way ANOVA with correction for multiple testing; details of the exact test used in each experiment are provided in the figure legends. Statistical analyses were carried out in GraphPad Prism version 10. In charts, statistical significance is represented as follows: n.s. – not significant; * - p<0.05; ** - p<0.01; *** - p<0.001; **** - p<0.0001.

## Acknowledgements

We thank Michael Ward for the i^3^N and i^3^N-CRISPRi lines and for helpful discussions. We thank the investigators mentioned in the Methods section for providing constructs.

