## Supplementary Materials for "Protrudin acts at ER-endosome contacts to promote KIF5-mediated endosomal fission and endosome-to-Golgi transport"

**Supplementary Figure Legends**

**Supplementary Figure 1. Validation of HeLa cells lacking protrudin. A**) HeLa cells were lentivirally transduced with a constructs containing the KRAB-deadCas9 (dCas9) coding sequence. After antibiotic selection, cells were lysed and immunoblotted against the antibodies shown. Tubulin immunoblotting serves as a loading control. **B**) CRISPRi HeLa cells expressing KRAB-dCas9 were transduced with one of 5 different single guide (SG) RNAs targeting *ZFYVE27* (the gene encoding protrudin) transcriptional start region. mRNA was extracted from the cells and RT-qPCR carried out to quantify *ZFYVE27* transcript expression in each line. *ACTB* was used as a housekeeping control to normalise the results. *ZYFVE27* mRNA expression is shown as percentage of expression in the scrambled control cell line. N = 1 biological repeat, each reaction was performed in triplicate. The lines SG1i and SG5i lines were used in subsequent experiments. **C-E**) Validation of the effect of protrudin depletion in the KRAB-dCas9 cells on the protrusion phenotype observed in spastin-depleted cells. **C**) HeLa-KRAB-dCas9 cells expressing a scrambled SG, SG1i or SG5i were transfected with a non-targeting siRNA or a pool of two siRNAs (1 and 3) targeting spastin, fixed and labelled with an antibody to acetylated tubulin. The mean +/- SEM percentage of cells with protrusions was quantified in (**D**), n = 3 biological repeats; 879 - 2038 cells per condition per repeat analysed. Statistical testing was done by repeated measures two-way ANOVA with Bonferroni correction for multiple comparisons; ns - not significant; *** - p<0.001. **E**) Spastin depletion in these experiments was confirmed by immunoblotting. GAPDH immunoblotting serves as a loading control.

**Supplementary Figure 2. SNX1 and VPS35 colocalise on endosomes and endosomal tubules. A**) Representative images of scrambled control and protrudin CRISPRi HeLa cells labelled for endogenous SNX1 and VPS35. The bottom three rows show higher magnification views of the boxed region in the corresponding panel in the upper row. Scale bars = 20 µm, inset scale bars = 2 µm. **B**) Quantification of colocalisation between SNX1 and VPS35 in scrambled control and protrudin CRISPRi HeLa cells. Pearson's colocalisation coefficient was measured for each cell using the ImageJ Coloc2 plugin. Mean +/- SEM of 3 biological repeats is plotted; 103 - 117 cells per condition per repeat analysed. Statistical testing used repeated measures one-way ANOVA with Dunnett correction for multiple comparisons; ns - not statistically significant.

**Supplementary Figure 3. Validation of depletion or inhibition of motor proteins in HeLa CRISPRi cell lines. A, B**) CRISPRi HeLa cells were lentivirally transduced with one of 3 different single guide (SG) RNAs targeting the KIF5B (**A**) or FYCO1 (**B**) transcriptional start region. mRNA was extracted from the cells and RT-qPCR carried out to quantify *ZFYVE27* transcript expression in each line. *ACTB* was used as a housekeeping control to normalise the results. *ZYFVE27* mRNA expression is shown as percentage of expression in the scrambled control cell line. n = 1 biological repeat, each reaction performed in triplicate. **C**) FYCO1 SG1i and SG3i cell lines were immunoblotted against endogenous FYCO1. GAPDH immunoblotting is shown to validate equal lane loading. **D**) Representative images of scrambled control and protrudin CRISPRi HeLa cells treated with 125 µM ciliobrevin D for 60 minutes or with vehicle control (DMSO). Following treatment, cells were fixed and labelled for the trans-Golgi marker TGN46. The characteristic Golgi dispersion seen after dynein inhibition was observed. Scale bar = 20 µm, inset scale bar = 2 µm.

**Supplementary Figure 4. $I^3$N validation. A** and **B**) $I^3$ iPSCs were differentiated to $I^3$Ns over a 21 day time course. Cells were lysed at the time points indicated (day 3, day 14 or day 21) and immunoblotted with the pluripotency (A; Nanog, Oct4, Sox2) or neuronal differentiation (B; MAP2, β-III tubulin or Tau) markers indicated. GAPDH blotting serves to validate equal lane loading. **C-F**) CRISPRi-$I^3$ iPSCs expressing SG RNAs targeting protrudin (**C** and **D**) or KIF5A (**E** and **F**) were differentiated to $I^3$Ns over a 14 day time course. RT-qPCR was carried out to quantify expression of the relevant transcript. *ACTB* was used as a housekeeping control to normalise the results and gene expression in the protrudin and KIF5A CRISPRi lines was normalised to scrambled control. n = 1 biological repeat, each reaction was performed in triplicate.

**Supplementary Movies 1 -3. SNX1 tubules in MRC5 cells.** CRISPRi GFP-SNX1 MRC5 cells expressing a scrambled SG (Movie 1), protrudin SG1i (Movie 2) or protruding SG5i (movie 3) were imaged by live cell microscopy. Movies were acquired at 2 frames per second and rendered at 30 frames per second.

Supplementary Movie 4. **mCherry-SNX1 and GFP-protrudin in MRC5 cells.** MRC5 cells stably expressing mCherry-SNX1 (green) and GFP-protrudin (purple) were visualised by live cell microscopy.  Movies were acquired at 2 frames per second and rendered at 30 frames per second.

### Supplementary Methods

#### siRNA transfection

Spastin and non-targeting control (NTC) small interfering RNAs (siRNAs) were obtained from Dharmacon and used at a concentration of 5 nM. To create double spastin and protrudin depletion using a mixed siRNA and CRISPRi approach, CRISPRi HeLa cells stably expressing protrudin SG1i and SG5i and control cells expressing scrambled SG1i were plated at a density of 2 x $10^4$ per well into six well plates. After 24 hours, cells were transfected with spastin or NTC siRNA using OligofectamineTM transfection reagent (ThermoFisher Scientific) according to the manufacturer's instructions, and harvested 72 hours later. SiRNA sequences were as follows:

Spastin siRNA 1 (D-014070-01) 5'-GAACUUCAACCUUCUAUAA

Spastin siRNA 3 (D-014070-03) 5'-UAUAAGUGCUGCAAGUUUA

NTC siRNA 1 (D-001210-01-05) 5'-UAGCGACUAAACACAACAA.

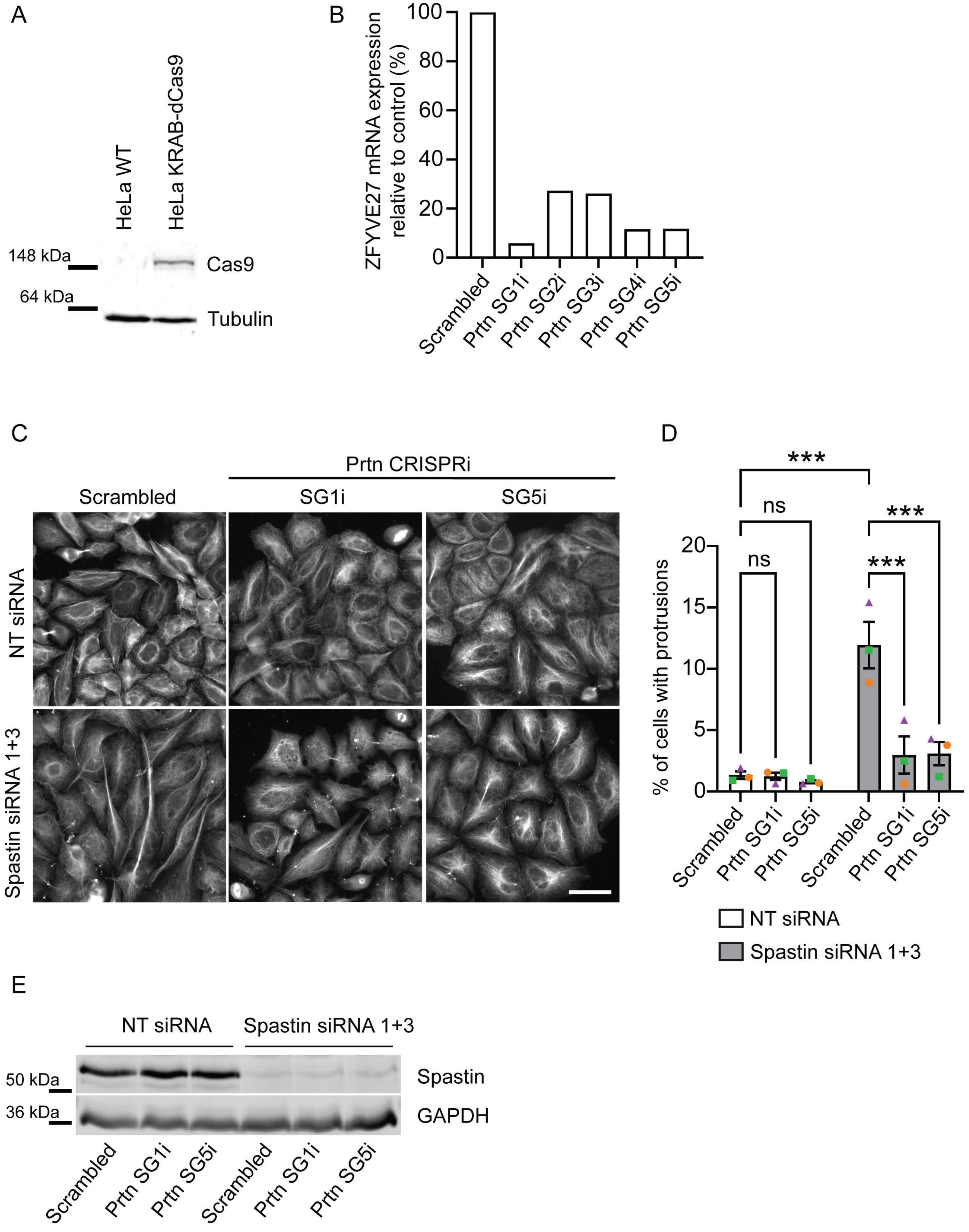


Supplementary Figure 1

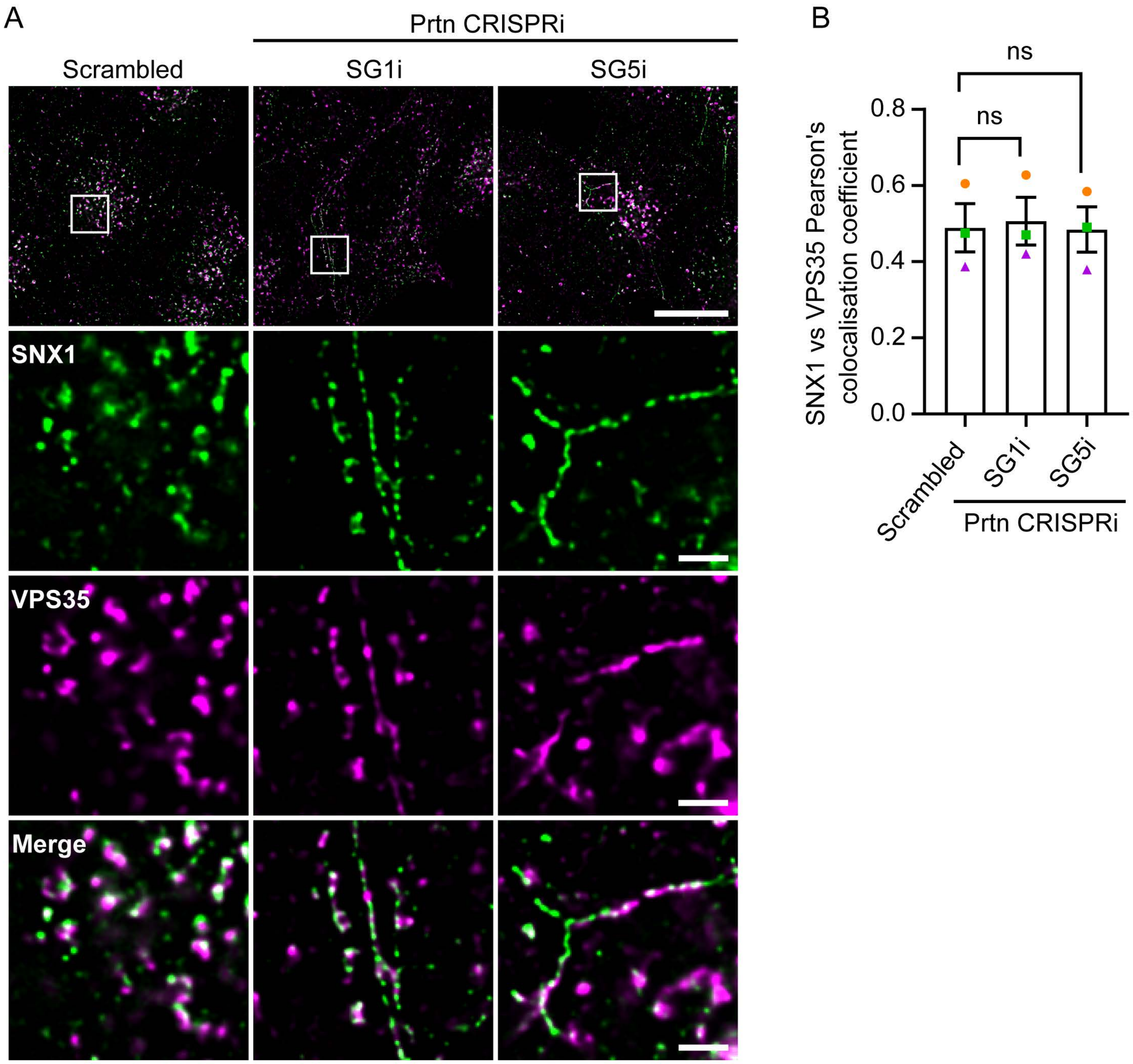


**Supplementary Figure 2**

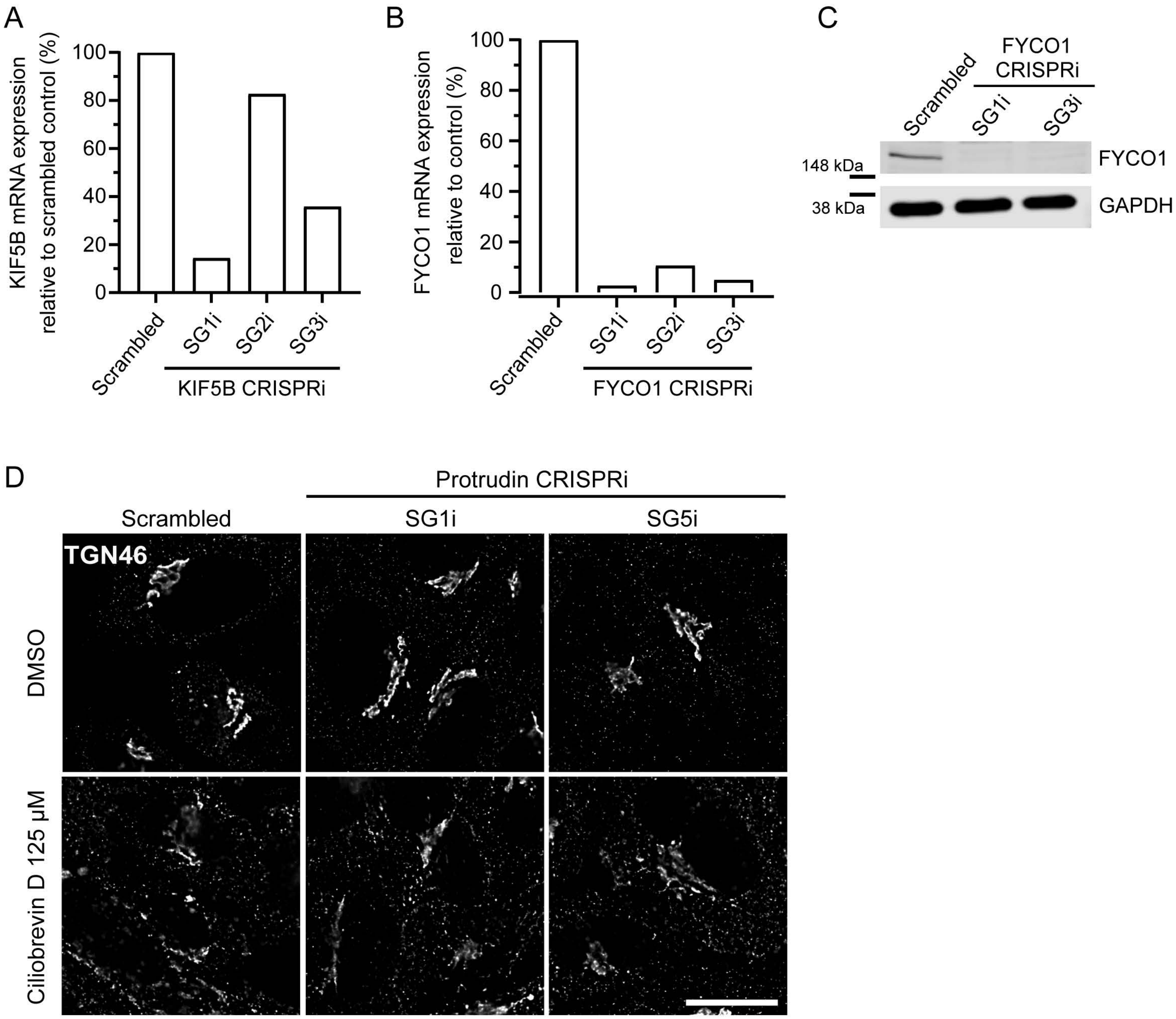


**Supplementary Figure 3**

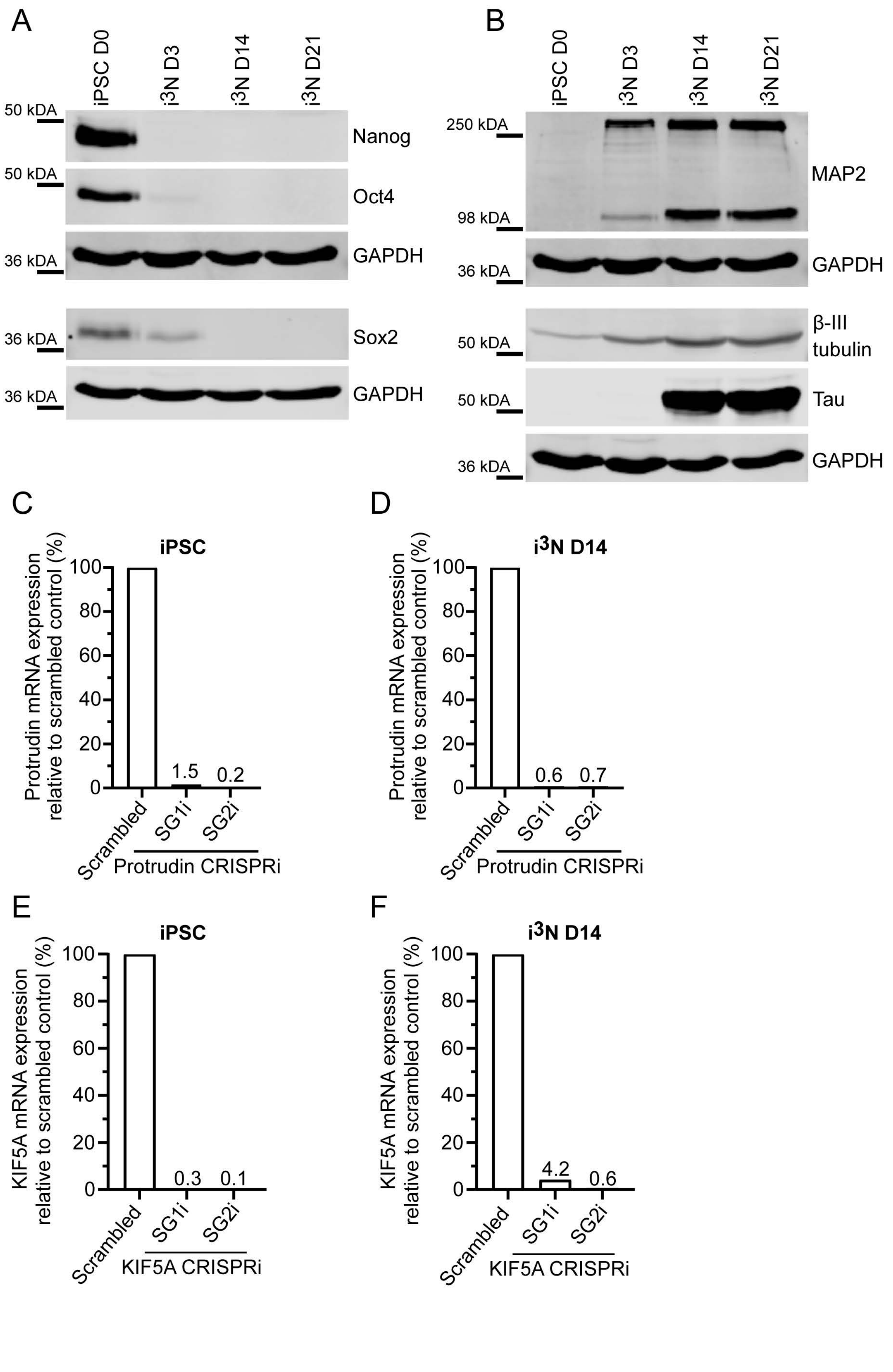


**Supplementary Figure 4**
